# Tuft cell-produced cysteinyl leukotrienes and IL-25 synergistically initiate lung type 2 inflammation

**DOI:** 10.1101/2021.09.26.461888

**Authors:** Saltanat Ualiyeva, Evan Lemire, Evelyn C. Aviles, Amelia A. Boyd, Caitlin Wong, Juying Lai, Tao Liu, Ichiro Matsumoto, Nora A. Barrett, Joshua A. Boyce, Adam L. Haber, Lora G. Bankova

## Abstract

Aeroallergen sensing by airway epithelial cells can trigger pathogenic immune responses leading to chronic type 2 inflammation, the hallmark of airway diseases such as asthma. Airway tuft cells are specialized chemosensory epithelial cells and the dominant source of the epithelial cytokine IL-25 in the trachea and of cysteinyl leukotrienes (CysLTs) in the naïve murine nasal mucosa. The interaction of IL-25 and CysLTs and the contribution of tuft cell-derived CysLTs to the development of allergen-triggered inflammation in the airways has not been clarified. Here we show that inhalation of LTC_4_ in combination with a subthreshold dose of IL-25 leads to dramatic synergistic induction of type 2 inflammation throughout the lungs, causing rapid eosinophilia, dendritic cell (DC) and inflammatory type 2 innate lymphoid cell (ILC2) expansion, and goblet cell metaplasia. While lung eosinophilia is dominantly mediated through the classical CysLT receptor CysLT_1_R, type 2 cytokines and activation of innate immune cells require signaling through both CysLT_1_R and CysLT_2_R. Tuft cell-specific deletion of the terminal enzyme requisite for CysLT production, *Ltc4s*, was sufficient to reduce both the innate immune response in the lung – eosinophilia, KLRG1^+^ ILC2 activation and DC recruitment – and the systemic immune response in the draining lymph nodes after inhalation of the mold aeroallergen *Alternaria*. Our findings identify surprisingly potent synergy of CysLTs and IL-25 downstream of aeroallergen-trigged activation of airway tuft cells leading to a highly polarized type 2 immune response and further implicate airway tuft cells as powerful modulators of type 2 immunity in the lungs.

**One Sentence Summary:** Tuft cells produce two highly synergistic mediators: LTC_4_ and IL-25 to cooperatively induce allergen-driven airway inflammation.

## INTRODUCTION

Type 2 immunity is a host defense mechanism engaged to expel helminths (*1*) and to repair epithelial cell damage from viral respiratory infections (*2*). Aeroallergens subvert this system through activation of epithelial cells for release of cytokines (IL-25, IL-33 and TSLP) and danger associated molecular patterns (DAMPs). DAMPs and epithelial cell cytokines act in concert to activate tissue resident dendritic cells (DCs), macrophages and innate type 2 lymphoid cells (ILC2s) to direct and propagate type 2 inflammation leading to chronic airway inflammatory diseases such as asthma and chronic rhinosinusitis (*3, 4*). Besides the classical epithelial cell cytokines, multiple recent studies suggest that DCs and ILC2s are also activated by lipid mediators (*5–7*) and neuropeptides (*8, 9*). While genetic and immunologic studies have defined barrier epithelial cell pathways that amplify mucosal inflammation, the events that initiate pathogenic immune recognition are still poorly understood.

Airway solitary chemosensory cells are unique epithelial cells that resemble taste bud cells in their expression of taste signaling molecules and have now been universally referred to as tuft cells (*10–12*). Unlike the chemosensory cells in taste buds, tuft cells are scattered as solitary cells in the epithelium of the upper and lower airway (*13*). The tuft cells of the airway are characterized by remarkable heterogeneity in morphology and receptor expression, leading to their definition as distinct cellular subsets (*11*). Tuft cells in the trachea are referred to as cholinergic brush cells (*14*), a similar population in the nasal respiratory mucosa is termed solitary chemosensory cells (SCCs) (*12*), and a much more abundant subset of chemosensory-like epithelial cells in the olfactory mucosa are referred to as microvillar cells (MVCs) (*15*). We recently demonstrated that tracheal brush cells, nasal respiratory SCCs and nasal olfactory MVCs share a core transcriptional profile with tuft cells from other tissue compartments – intestinal, thymic and gallbladder tuft cells – suggesting they belong to one large family of chemosensory tuft cells with tissue specific differences mostly in bitter and sweet taste receptor expression (*16–18*). Among the shared markers of all airway tuft cells from the nasal olfactory, respiratory and tracheal mucosa are TRPM5, required for Ca^2+^ triggered signal transduction in sweet and bitter taste receptor expressing cells (*12, 13, 15, 16, 19*), the epithelial cytokine IL-25 (*20, 21*), the enzyme choline acetyltransferase (ChAT) for acetylcholine generation (*14*) and the transcription factor *Pou2f3* (*17, 18, 22*). Notably, a core transcriptional feature of all tuft cells are the transcripts encoding the lipoxygenase (*Alox5, Alox5ap* and *Ltc4s*) and cyclooxygenase (*Ptgs1* and *Hpgds*) pathway enzymes (*11, 16, 23, 24*). We recently reported the functional capacity of tuft cells to generate high levels of cysteinyl leukotrienes (CysLTs), the major products of the lipoxygenase pathway (*16*), *ex vivo* at a level exceeding that of naïve hematopoietic cells in the naïve nasal mucosa. Aeroallergens like the mold aeroallergen *Alternaria* and the dust mite allergen *Dermatophagoides pteronyssinus* trigger a P2y2 receptor-dependent autocrine loop leading to generation of CysLTs from tuft cells (*16*). Although we had demonstrated that airway tuft cells can generate CysLTs in response to allergens, how tuft cell-derived CysLTs regulate allergen-triggered type 2 immunity in the airways has not been defined.

CysLTs are named for their canonical generation by leukocytes recruited or activated in the setting of established inflammation. Following receptor-mediated Ca^2+^ flux, phospholipase A2 (PLA2α) releases phospholipids at the outer nuclear membrane to generate free arachidonic acid. Arachidonic acid is metabolized by the cyclooxygenase and lipoxygenase pathways to prostaglandins and cysteinyl leukotrienes, respectively. The first enzyme of the lipoxygenase pathway, 5-lipoxygenase (5-LO), oxidizes arachidonic acid in the presence of 5-LO activating protein (FLAP) to generate leukotriene A_4_ (LTA_4_), which is subsequently converted to LTC_4_ by leukotriene C_4_ synthase (LTC_4_S) (*25*). LTC_4_ is rapidly exported by the ATP-dependent multidrug resistance protein 1 (MRP1) and converted within minutes to LTD_4,_ which in turn is rapidly metabolized to LTE_4_, a more stable but less potent terminal CysLT product (*26*). CysLTs exert their effects through three G-protein coupled receptors. CysLT_1_R has high affinity for LTD_4_ (Kd ∼ 1 nM) and binds LTC_4_ with lesser affinity, and can transduce signals to LTE_4_ as well (*27*). CysLT_2_R, which is resistant to currently available pharmacotherapy, targeting CysLT_1_R binds LTC_4_ and LTD_4_ at equimolar concentrations (*28*). CysLT_3_R (OXGR1 or GPR99) is a high affinity receptor for LTE_4_ (*29*). CysLTs potently augment established type 2 lung inflammation (*30, 31*). This might be at least partially explained by the ability of CysLTs to potentiate the effects of epithelial cytokines. LTC_4_ synergizes with IL-33 for ILC2 activation in the airways and associated lung eosinophilia through CysLT_1_R-mediated NFATc translocation (*5–7*). LTC_4_ also has CysLT_1_R-mediated additive effects on IL-25-induced activation of intestinal ILC2s *in vitro* (*32*). The fact that tuft cells generate both LTC_4_ and IL-25 suggests the potential for these two mediators to facilitate type 2 airway inflammation *in vivo*, but this possibility has not yet been explored.

Here, we demonstrate that two mediators generated by tuft cells – LTC_4_ and IL-25 – synergize to drive airway type 2 inflammation. Exogenous LTC_4_ or low dose IL-25 alone only weakly induced inflammation, but the combination of IL-25 and LTC_4_ administered intranasally strongly induced lung type 2 cytokine expression, as well as lung ILC2 and DC expansion and airway eosinophilia. The proinflammatory synergistic effect of LTC_4_ and IL-25 is reduced in mice lacking the two receptors for LTC_4_ – *Cysltr1^-/-^* and *Cysltr2^-/-^* – with a dominant effect of CysLT_1_R in regulating eosinophil recruitment. Tuft cell-targeted deletion of the terminal CysLT biosynthetic enzyme, *Ltc4s*, in *Chat^Cre^Ltc4s^fl/fl^* mice led to a specific reduction of CysLT generation in tuft cells while preserving their homeostatic transcriptional profile and leaving other CysLT sources unaffected. While *Ltc4s* deletion ablated the ability of tuft cells to generate CysLTs, it did not diminish their ability to respond to ATP with generation of PGD_2_, a cyclooxygenase pathway-derived eicosanoid that is also involved in type 2 inflammation. The early lung eosinophilia and the associated lung draining lymph node hyperplasia triggered by the inhalation of the mold aeroallergen *Alternaria* was reduced in *Chat^Cre^Ltc4s^fl/fl^* mice, as was ILC2 activation and expansion in the lung and lung DC expansion. Thus, tuft cells are unique epithelial effector cells capable of inducing eosinophilic type 2 lung inflammation through synergistic cytokine and lipid mediator signaling.

## RESULTS

### Exogenous LTC_4_ dramatically amplifies IL-25 induced lung inflammation

The role of tuft cells as initiators of type 2 inflammation was initially attributed to their production of IL-25 (*20, 21, 33*). Our work identified LTC_4_ as a major product of airway tuft cells after aeroallergen activation. We therefore investigated the behavior of these pro-inflammatory mediators *in vivo.* We administered LTC_4_ alone at a concentration of 1.6 mmol, which induces only mild inflammation in naive mice, or low dose IL-25 alone (100 ng or 5 times lower than the dose used for most airway inflammation studies (*9*)) or a combination of IL-25 and LTC_4_ to wild-type (WT) mice (**Fig. 1A**). Intranasal application of low-dose IL-25 or high dose LTC_4_ induced very mild lung inflammation, while the combination of LTC_4_ and IL-25 led to an increase of CD45^+^ cells in the lung (**Fig. 1B**). The lung infiltrate was dominated by eosinophils, which accounted for 33% of the hematopoietic cells by fluorescence-activated sorting (FACS) (**Fig. 1C, D**) and were the most abundant inflammatory cells in the perivascular infiltrate in the lung (**S1A, B**). When we assessed the ILC2 compartment (**Fig. S1C**) for changes in response to IL-25 and LTC_4_, we detected a population of KLRG1^+^ ICOS^+^ Thy1.2^+^ ILC2s, similar to the previously described inflammatory ILC2s (*34*) (**Fig. 1E**). Interestingly, inhaled low dose IL-25 was insufficient to induce this ILC2 subset, but the addition of LTC_4_ led to a 30% increase compared to IL-25 or LTC_4_ alone (**Fig. 1E-G**). This expansion was specific to inflammatory ILC2s, as we did not detect a significant change in the number of the unfractionated lin^-^Thy1.2^+^ ILC2s or any of the subsets of KLRG1^-^ILC2s (**Fig. 1H and Fig. S1D-F**). The IL-25/LTC_4_ synergistic increase in lung eosinophilia and inflammatory ILC2 expansion was associated with an expansion of pulmonary DCs (**Fig. S1G**), particularly the CD301b^+^ subset (**Fig. 1I and Fig. S1H**). Despite the clear induction of lung inflammation, the number of tuft cells assessed in whole tracheal mounts did not change (**Fig. 1J**), unlike the effect of LTE_4_ on tracheal tuft cells (*20*). This suggests that conversion to LTE_4_ was not responsible for the observed effects of exogenous LTC_4_ *in vivo*. Conversely, lung remodeling with significant goblet cell metaplasia was strongly induced by the combination of LTC_4_ and IL-25 (**Fig. 1JK, L**). Consistent with the effect on inflammatory cell recruitment and goblet cell metaplasia, LTC_4_ + IL-25 augmented type 2 cytokine production in the naïve mouse lung over PBS or either mediator alone, with significant increases of three type 2 cytokines – IL-4, IL-5 and IL-13 (**Fig. 1M**). In addition, we detected a significant increase in lung IL-6 and a small but significant augmentation of TNF-α (**Fig. S2A, B**). The type 1 cytokine IFN-γ was undetectable and IL-17 family member cytokines were not induced (**Fig. S2**).

**Fig. 1.**
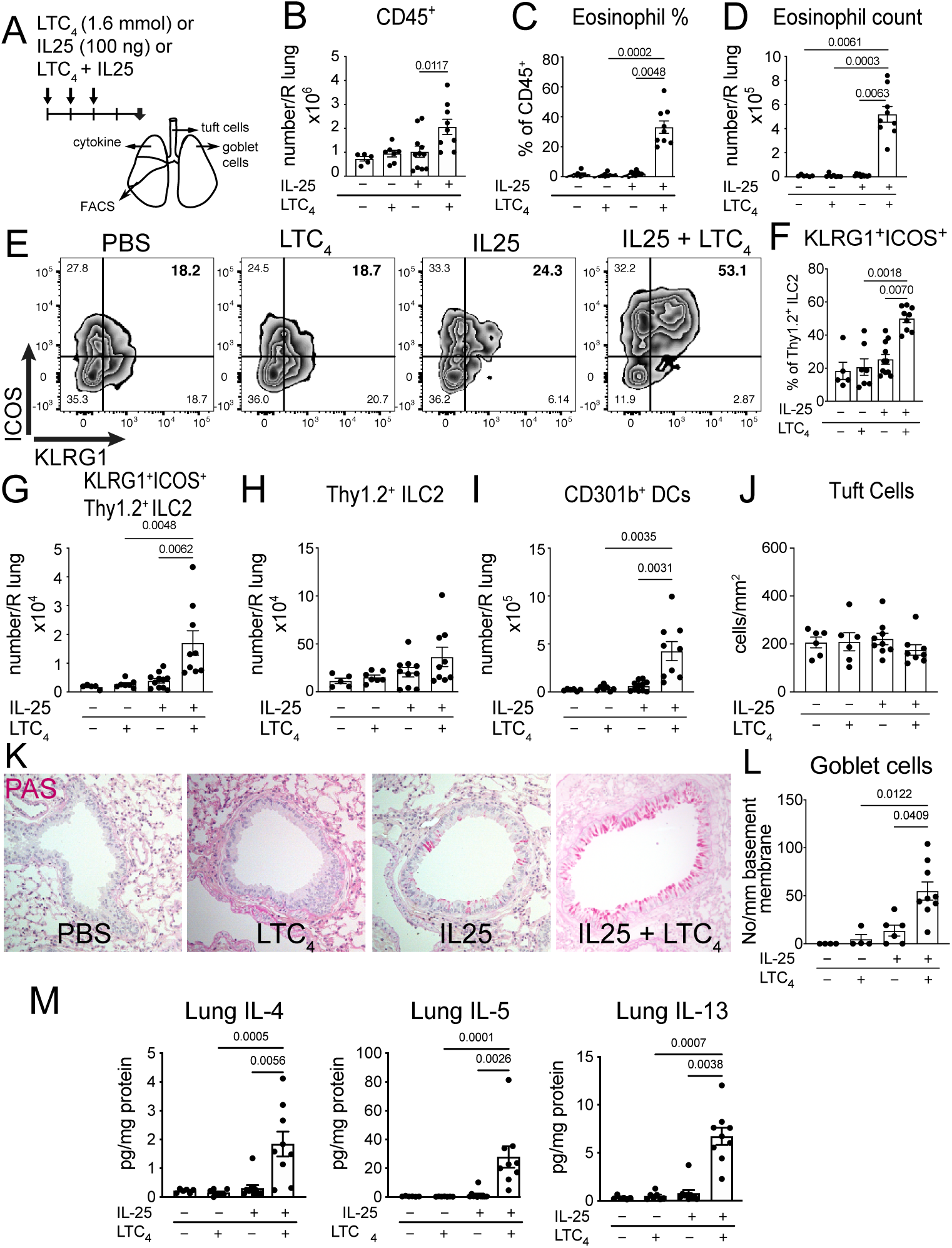
LTC_4_ and IL-25 synergize for airway type 2 lung inflammation. (**A**) WT mice were given three daily inhalations of LTC_4_ (1.6 mmol), or IL-25 (100 ng) or a combination of LTC_4_ and IL-25 and assessed 2 days after the last dose. The number of lung CD45^+^ cells (**B**), frequency and number of eosinophils (**C, D**) frequency and number of KLRG1^+^ICOS^+^Thy1.2^+^ ILC2s (**F, G**), number of unfractionated Thy1.2^+^ ILC2s (**H**) and CD301b^+^MHCII^+^CD11b^+^ DCs (**I**) were assessed by FACS. (**E**) Representative FACS plots of KLRG1 and ICOS expression in lin^-^Thy1.2^+^ ILC2s. (**J**) The number of tuft cells was assessed in whole tracheal mounts. (**K**) Goblet cell metaplasia was assessed in cross sections of the lung with Periodic acid Schiff (PAS) stain and goblet cells were enumerated per mm of basement membrane in the large bronchi (**L**). (**M**) Lung cytokine concentration was determined by LegendPlex and calculated per mg of lung protein extracted. Data are means ± SEM pooled from 3 independent experiments, each dot is a separate mouse, Kruskal-Wallis ANOVA with Dunn’s correction for multiple comparisons, p values <0.05 indicated.

Together, this data demonstrates that two highly synergistic pro-inflammatory and airway remodeling pathways can be triggered by the tuft cell mediators IL-25 and LTC_4_, and that the presence of LTC_4_ dramatically reduces the IL-25 dose required to induce type 2 inflammation. Furthermore, tuft cell mediators act in synergy to effectively skew innate inflammation to a type 2 phenotype.

### The synergy of LTC_4_ and IL-25 is cooperatively mediated by CysLT_1_R and CysLT_2_R

To define the CysLT receptors responsible for mediating the synergy of IL-25 and LTC_4_, we evaluated the response in respective strains of mice lacking the three CysLT receptors. This allowed us to account for both the direct effect of LTC_4_ on CysLT_1_R and CysLT_2_R and of the biosynthetic products LTD_4_ and LTE_4_. We found a near complete abrogation of lung inflammation and eosinophil recruitment in *Cysltr1^-/-^* mice (**Fig. 2A, B**) with marginal non-significant reduction of eosinophil counts in *Cysltr2^-/-^* or *Cysltr3^-/-^* mice (**Fig. 2A, B**). Neither CysLT_1_R nor CysLT_2_R deletion abolished the specific increase in the percent of KLRG1^+^ICOS^+^Thy1.2^+^ILC2 cells induced by LTC_4_ + IL-25, which suggests a cooperative effect with the IL-25 receptor IL17RB (**Fig. 2C, D**). While the percent of KLRG1^+^ICOS^+^Thy1.2^+^ILC2 did not change, the absolute number of these inflammatory ILC2s was reduced in both *Cysltr1^-/-^* and *Cysltr2^-/-^* mice (**Fig. 2E**). Thus, although IL17RB signaling is likely a driver of the expansion of inflammatory ILC2s (*34*), co-signaling with CysLT receptors can trigger the inflammatory ILC2s even at low levels of IL-25. Inflammatory CD301b^+^ DCs were reduced in *Cysltr1^-/-^*, with a non-significant trend toward reduction in *Cysltr2^-/-^* mice (**Fig. 2F**). Finally, type 2 cytokines were significantly reduced in both *Cystlr1^-/-^* and *Cystlr2^-/-^* mice, with a predominant reduction in *Cystlr1^-/-^* mice (**Fig. 2G**). Collectively, our data highlights the complexity of the CysLT system and implicates two receptors in the effects of LTC_4_ on target cells. While CysLT_1_R deletion completely abrogates the synergistic effect of LTC_4_ and IL-25 on lung eosinophil recruitment and partially reduces the expansion of ILC2s, deleting both LTC_4_ receptors and the IL-25 receptor IL17RB is likely required to completely abolish the effect of the potent LTC_4_/IL-25 synergy.

**Fig. 2.**
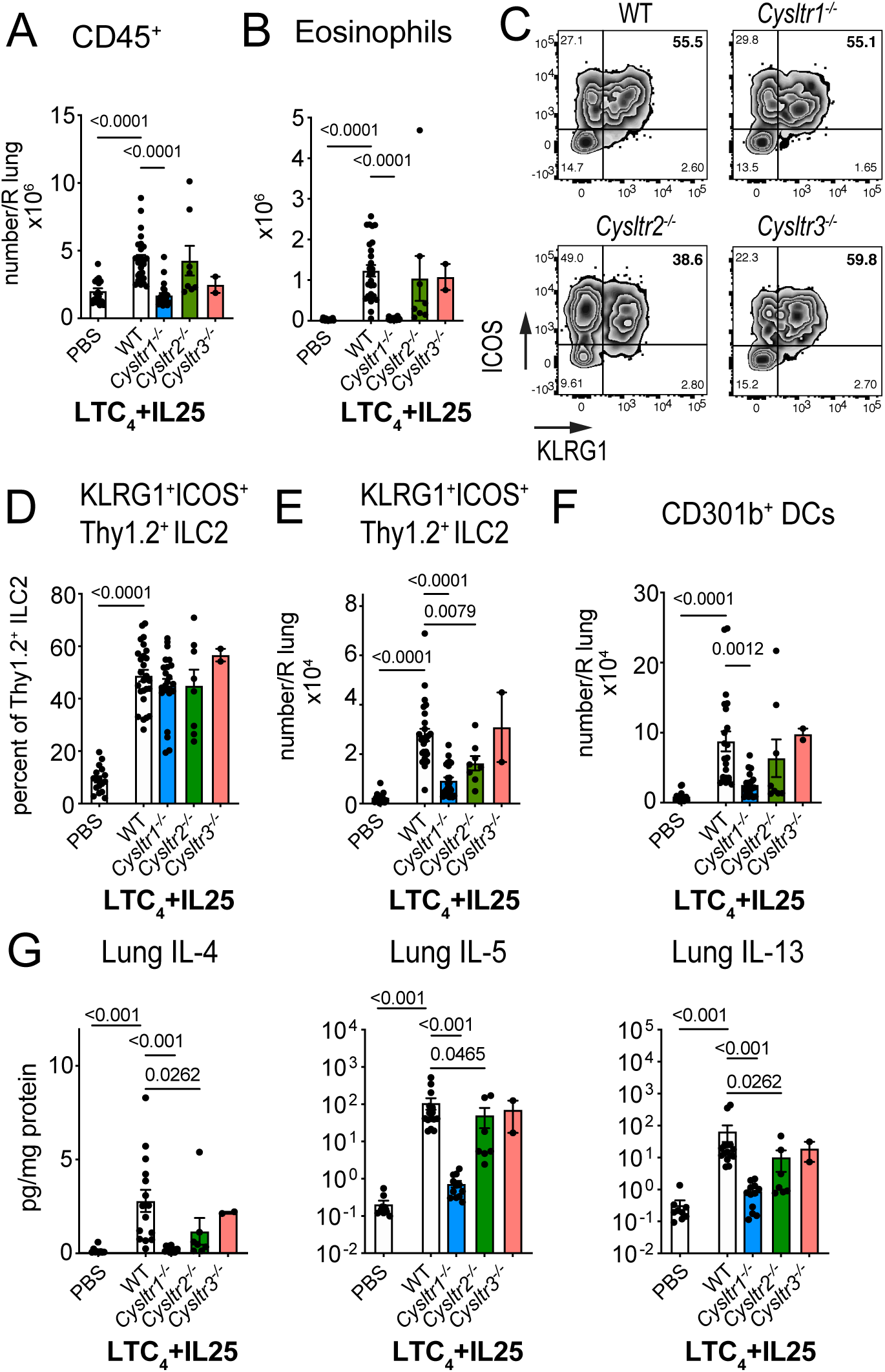
LTC_4_ potentiation of IL-25-induced type 2 inflammation depends on CysLT_1_R and CysLT_2_R. WT (C57BL/6), *Cysltr1^-/-^* , *Cysltr2^-/-^* and *Cysltr3^-/-^* mice were given three daily inhalations of LTC_4_ and IL-25 daily for 3 days and assessed 2 days after the last dose. Numbers of lung CD45^+^ cells (**A**), eosinophils (**B**), frequency and number of KLRG1^+^ICOS^+^ ILC2s (**D, E**), and number of CD301b^+^ CD11b^+^ DCs (**F**) were assessed by FACS. (**C**) Representative FACS plots of KLRG1 and ICOS expression in lin^-^Thy1.2^+^ ILC2s. (**G**) Cytokine protein concentration was determined by LegendPlex and expressed per mg of lung protein. Data are means ± SEM pooled from 3 independent experiments, each dot is a mouse, significant p values indicated for each pairwise comparison: Mann Whitney U test.

### ChAT-mediated targeted deletion of *Ltc4s* selectively ablates tuft cell generation of CysLTs

To assess the epithelial and immune effector circuit regulated by tuft cell-derived CysLTs, we generated *Ltc4s^fl/fl^* mice to enable conditional deletion of *Ltc4s*. We first characterized the populations of epithelial cells in the nose in the *Ltc4s^fl/fl^* mice and compared them to nasal tuft cells and epithelial cells from ChAT-eGFP mice. In the absence of a fluorescent marker in the *Ltc4s^fl/fl^* mice, we used our previously published combination of EpCAM and CD45 expression to isolate tuft cell-enriched populations from the nasal mucosa (*16*). We isolated EpCAM^high^CD45^low^ cells to compare them to EpCAM^high^CD45^-^ and EpCAM^interm^ cells from the same *Ltc4s^fl/fl^* mice (**Fig. S3A**). We then compared these three populations derived from the *Ltc4s^fl/fl^* mice to the more pure populations of tuft cells derived from the olfactory and respiratory mucosa of ChAT-eGFP mice. In ChAT-eGFP mice, we had characterized two populations of nasal tuft cells based on their FACS morphology, which share the core transcriptional profile of tuft cells from other mucosal organs, but differ in the expression of taste receptors and roughly correspond to olfactory MVCs and respiratory SCCs (*16*). Here, we isolated ChAT-eGFP SSC^low^ tuft cells from the olfactory mucosa and ChAT-eGFP SSC^high^ tuft cells from the respiratory mucosa to more clearly delineate the MVCs and SCCs (**Fig. S3B**). We confirmed that the transcriptional profile of EpCAM^high^CD45^low^ epithelial cells is highly similar to the dominant nasal population of olfactory tuft cells (**Fig. 3A and S3C, D**). Furthermore, EpCAM^high^CD45^low^ cells share the core transcriptional profile of tracheal tuft cells defined by scRNAseq (*11*) while EpCAM^high^CD45^-^ likely represent a less pure mix of tuft cells and epithelial cells (**Fig. 3B**). Both EpCAM^high^ populations strongly expressed the tuft cell-specific genes *Trpm5, Avil,* and *Pou2f3* as well as enzymes in the CysLT biosynthetic cascade, including *Alox5, Alox5ap* and *Ltc4s* (**Fig. 3C, D**), with a higher level of all transcripts found in the EpCAM^high^ CD45^low^ cells.

**Fig. 3.**
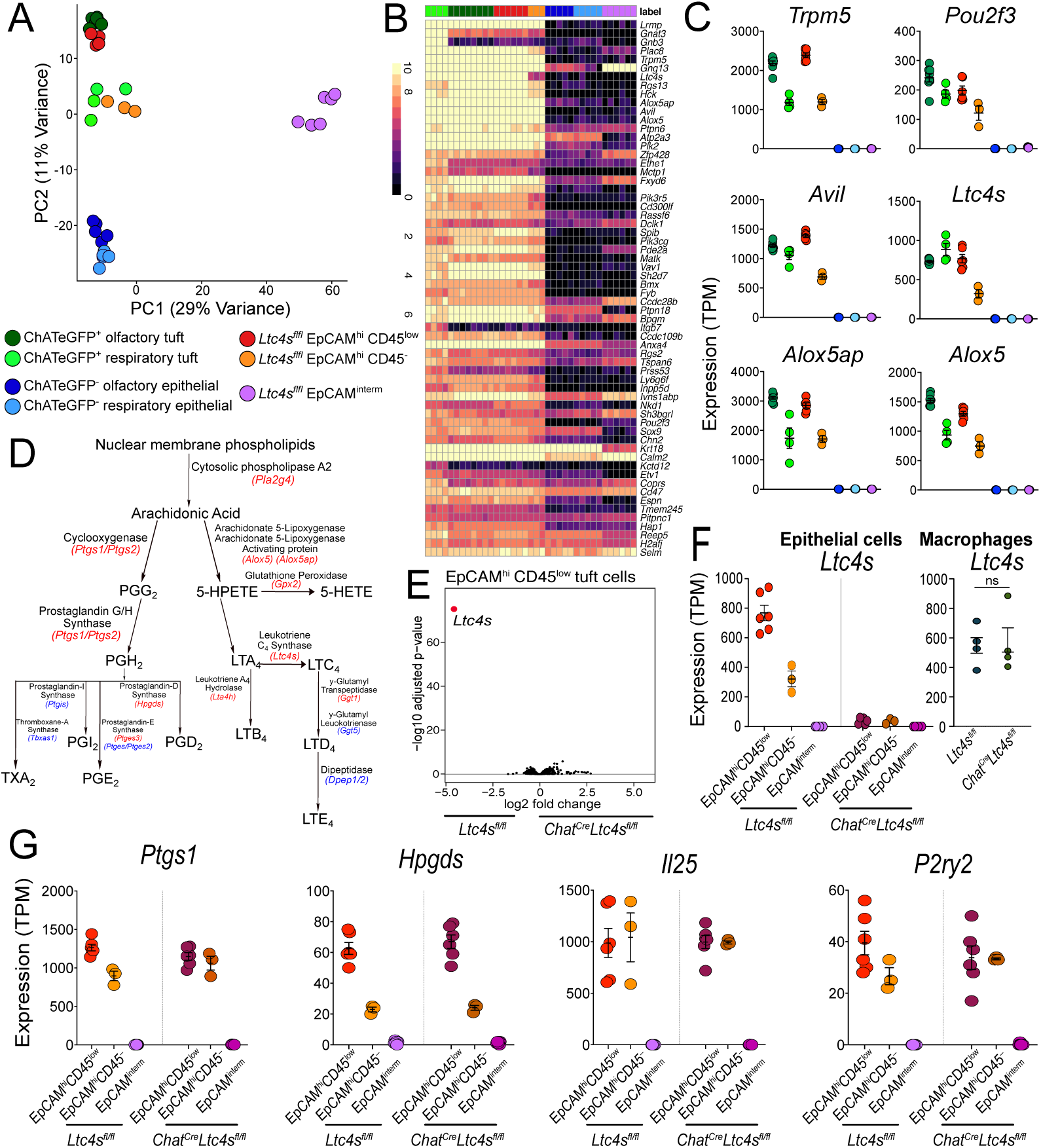
Essential features of the tuft cell transcriptome are preserved after specific deletion of *Ltc4s*. Tuft cells and epithelial cells were isolated from the naïve nasal mucosa of ChAT-eGFP, *Ltc4s^fl/fl^* and *ChAT^Cre^Ltc4s^fl/fl^* mice. Tuft cells were defined as EpCAM^high^GFP^+^SSC^low^ olfactory tuft and EpCAM^high^GFP^+^SSC^high^ respiratory tuft in ChAT-eGFP mice and compared with EpCAM^high^CD45^low^ and EpCAM^high^CD45^-^ and EpCAM^interm^ epithelial cells from *Ltc4s^fl/fl^*. (**A**) Principal components analysis comparing each of the above populations of cells from ChAT-eGFP mice and control *Ltc4s^fl/fl^* mice. Numbers indicate frequency of transcripts described by each principal component (PC). (**B**) Expression of the tracheal scRNAseq signature (*11*) genes across tuft and non-tuft epithelial cells in the nose. (**C**) TPM counts of the indicated genes derived from RNA seq analysis using DeSeq2 showing tuft cell specific genes. (**D**) Eicosanoid pathway enzymes and transcripts (in italics). Highlighted in red are transcripts expressed in tuft cells in RNAseq datasets, and in blue, transcripts that were not detected. (**E**) Volcano plot displaying significance (y axis, -log10(FDR)) of differential expression (x axis, log2 fold-change) in EpCAM^high^CD45^low^ tuft cell-enriched cells between *Ltc4s^fl/fl^* and *Chat^Cre^Ltc4s^fl/fl^* mice. (**F**) *Ltc4s* expression in with EpCAM^high^CD45^low^ and EpCAM^high^CD45^-^ and EpCAM^interm^ epithelial cells and macrophages derived from *Ltc4s^fl/fl^* and *Chat^Cre^Ltc4s^fl/fl^* mice, ns = not significant (**G**) Expression level (TPM) of the indicated genes derived from RNA seq analysis using DeSeq2 of EpCAM^+^ cells from *Ltc4s^fl/fl^* and *Chat^Cre^Ltc4s^fl/fl^* mice.

We then crossed the *Ltc4s^flfl^* mice to *Chat^Cre^* mice to specifically delete *Ltc4s* in ChAT-expressing cells. Classically, cholinergic neurons are the best characterized *Chat*-expressing cells (*35, 36*). Therefore, to account for possible *Chat* and *Ltc4s* coexpression in the central and peripheral nervous system, we interrogated published scRNAseq datasets of the mouse central and peripheral nervous system and detected no expression of *Ltc4s* in the main *Chat*-expressing populations of the nervous system – motor neurons and visceral motor neurons (**Fig. S4A**). In the brain tissue, *Ltc4s* was only expressed in immune cell populations. *Chat* expression in immune cell subsets has been reported in CD8 and CD4 T cells (*37*) and more recently in ILC2s (*38, 39*), while among hematopoietic cells in the nasal mucosa, *Ltc4s* is reported in macrophages, DCs, and mast cells (*40, 41*). To assess for co-expression of *Chat* and *Ltc4s* more broadly in the airway mucosa, we evaluated a recently generated dataset of 50,000 epithelial and immune cells in the nasal mucosa (derived from a recently collected unpublished scRNAseq dataset from our laboratory of epithelial and immune cells from naïve and *Alternaria-*challenged mice). Several immune cell subsets – including T and B cell subsets, ILC2s and macrophages – expressed *Chat*, while macrophages, eosinophils, and DCs expressed *Ltc4s* as expected (**Fig. S4B**). Remarkably, only the tuft cells expressed both *Chat* and *Ltc4s* in > 10% of the cells (**Fig. S4B**). Consistent with this co-expression, ChAT^Cre^ mediated deletion of *Ltc4s* led to a profound 95% reduction of *Ltc4s* in the EpCAM^high^CD45^low^ tuft cell-enriched population and a 90% reduction in the EPCAM^high^CD45^-^ cells (**Fig. 3E, F**). To confirm that *Ltc4s* expression was not affected in macrophages and DCs, we isolated both immune cell subsets from *Ltc4s^flfl^* and *Chat^Cre^Ltc4s^flfl^* mice (**Fig. S4C**). Macrophages expressed *Ltc4s* at similarly high levels as tuft cells with no reduction in *Ltc4s* expression in *Chat^Cre^Ltc4s^flfl^* mice (**Fig. 3F**). DC expression of *Ltc4s* was magnitudes lower than that of macrophages and tuft cells and was not significantly affected in *Chat^Cre^Ltc4s^flfl^* mice (**Fig. S4D**). Thus, ChAT^Cre^ mediated deletion of *Ltc4s* effectively ablated expression of the *Ltc4s* transcript in tuft cells without an effect on other epithelial, neuronal or immune cell subtypes.

To probe for homeostatic effects of *Ltc4s* deletion on tuft cells, we compared the transcriptional profile of EpCAM^high^CD45^low^ tuft cells from *Ltc4s^fl/fl^* and *Chat^Cre^Ltc4s^fl/fl^* mice. While *Ltc4s* expression was 95% reduced in the tuft cell-enriched EpCAM^high^CD45^+^ cells of *Chat^Cre^Ltc4s^fl/fl^* mice, we detected a very limited effect on the homeostatic transcriptional profile of this tuft cell-enriched population besides *Ltc4s* (**Fig. 3E**). EpCAM^high^CD45^low^ tuft cells from *Chat^Cre^Ltc4s^fl/fl^* mice clustered together with EpCAM^high^CD45^low^ tuft cells from *Ltc4s^fl/fl^* mice, suggesting no significant transcriptional changes (**Fig. S5A, B**). In addition, tuft cell numbers in EpCAM^high^CD45^low^ cells in the nose did not differ significantly between *Ltc4s^fl/fl^* and *Chat^Cre^Ltc4s^fl/fl^* mice (**Fig. S5C**). Further, deletion of *Ltc4s* in tuft cells did not affect the homeostatic expression of other eicosanoid biosynthetic enzymes in tuft cells (**Fig. 3G and Fig. S5D**). Finally, expression of the other major effector mediator generated by tuft cells, *Il25*, and the expression of P2Y2R, involved in allergen recognition by tuft cells (*16*), were not reduced by the tuft cell-specific deletion of *Ltc4s* (**Fig. 3G**). Collectively, we found that ChAT^Cre^-mediated deletion of *Ltc4s* in tuft cells specifically targets *Ltc4s* with minimal effect on the tuft cell homeostatic transcriptional profile.

To confirm that the *Ltc4s* transcript deletion leads to a specific impairment of CysLT generation, we isolated nasal EpCAM^high^SSC^low^ epithelial cells from *Chat^Cre^, Ltc4s^fl/fl^* and *Chat^Cre^Ltc4s^fl/fl^* mice and assessed their responses to Ca^2+^ ionophore and the physiologic tuft cell stimulus ATP using an *ex vivo* stimulation assay (*16*). Tuft cells isolated from both *Chat^Cre^* and *Ltc4s^fl/fl^* mice generated high levels of CysLTs in response to Ca^2+^ ionophore, and this response was almost entirely absent in tuft cells from *Chat^Cre^Ltc4s^fl/fl^* mice (**Fig. 4A**). CD45^+^ cells isolated from *Ltc4s^fl/fl^* and *Chat^Cre^Ltc4s^fl/fl^* mice stimulated with Ca^2+^ ionophore generated CysLTs at comparable levels, confirming the specificity of *Ltc4s* deletion to tuft cells (**Fig. 4B**). Stimulation with ATP, a physiologic stimulus of tuft cells through the P2Y2 receptor (*16*), similarly elicited CysLTs from tuft cells isolated from *Ltc4s^fl/fl^* but not from *Chat^Cre^Ltc4s^fl/fl^* mice (**Fig. 4C**). Consistent with the RNAseq data demonstrating preserved cyclooxygenase biosynthetic transcripts, *Chat^Cre^Ltc4s^fl/fl^* derived tuft cells stimulated with both the Ca^2+^ ionophore A23187 and with ATP generated PGD_2_ at levels comparable to tuft cells sorted from *Ltc4s^fl/fl^* and *Chat^Cre^* mice (**Fig. 4D, E**). We could not verify the effect of CysLT deletion on IL-25 production at the protein level, but confirmed that IL-25 transcript was not reduced in these *Chat^Cre^Ltc4s^fl/fl^* mice (**Fig. 3G**). We had previously shown that tuft cells are the dominant source of *Alternaria-* triggered nasal CysLTs *in vivo* using mice with deletion of the tuft cell-specific transcription factor *Pou2f3* (*16*). Here, intranasal *Alternaria* administered *in vivo* to *Ltc4s^fl/fl^* mice induced generation and luminal release of CysLTs into the nasal lavage fluid, which was reduced in *Chat^Cre^Ltc4s^fl/fl^* mice, confirming that *Ltc4s* expressed in tuft cells is a critical contributor to the innate nasal LTC_4_-generating response to aeroallergens (**Fig. 4F**).

**Fig. 4.**
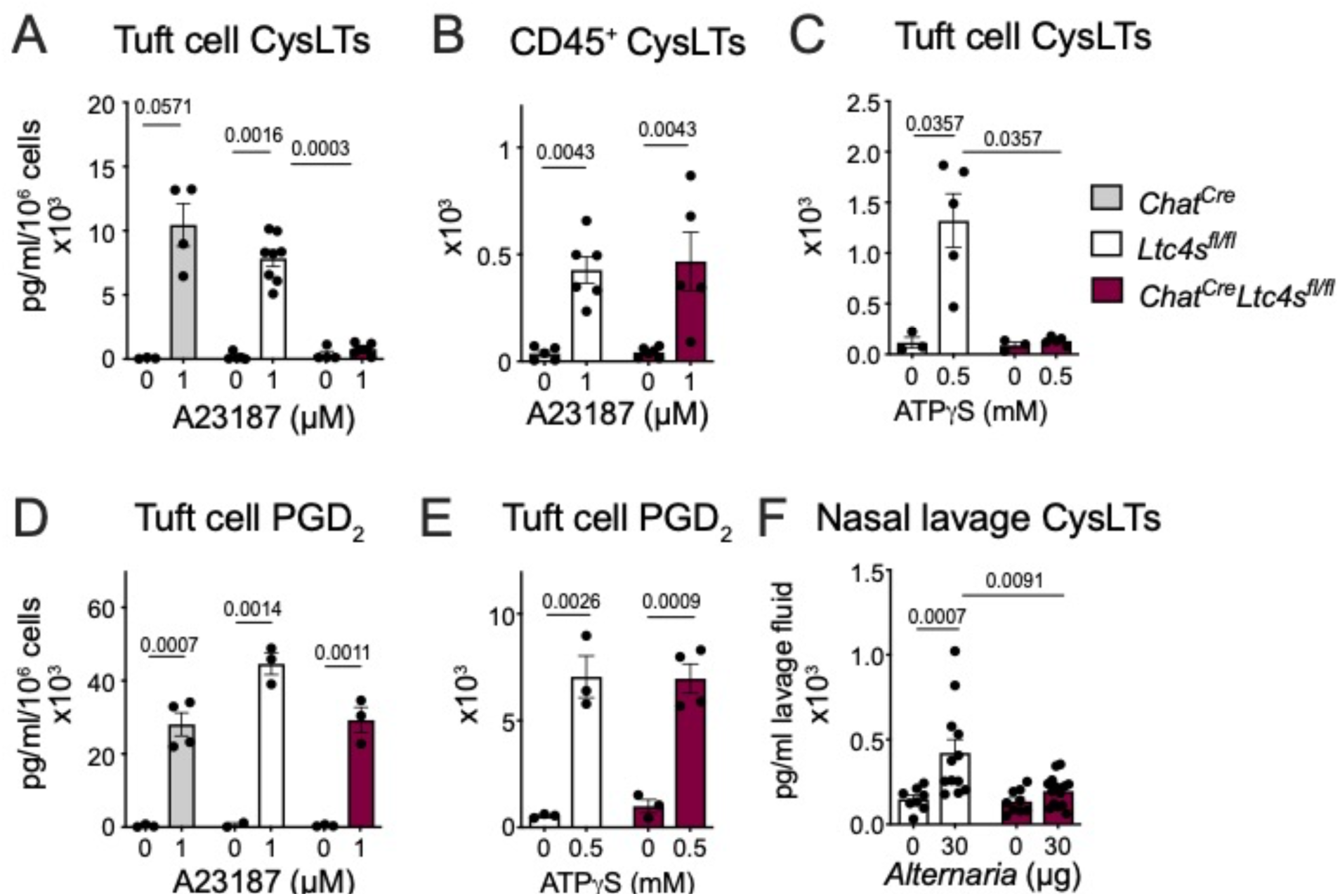
Tuft cell-specific *Ltc4s* deletion does not alter their ability to respond to activating signals. (**A-E**) EpCAM^high^CD45^low^ SSC^low^ tuft cells (**A, C-E**) or CD45^+^ cells (**B**) were isolated from the nasal mucosa of *Chat^Cre^, Ltc4s^flfl^* and *Chat^Cre^Ltc4s^flfl^* mice and stimulated *ex vivo* with Ca^2+^ ionophore (A23187) (**A, B, D**) or ATPγS (**C, E**). The concentration of CysLTs (**A-C**) or PGD_2_ (**D, E**) in the supernatants was measured by ELISA at 30 min. (**F**) *Alternaria* was administered intranasally to naïve *Ltc4s^flfl^* and *Chat^Cre^ Ltc4s^flfl^*mice and nasal lavage (NL) was obtained at 30 min. The concentration of CysLTs (**F**) were measured by ELISA after acetone precipitation. Data are means ± SEM, from at least three independent experiments, each dot represents a separate biological replicate representing pooled cells from 1-3 mice, Mann Whitney U-test p values <0.05 are indicated.

Collectively, these data demonstrate that *Chat^Cre^-*mediated deletion of *Ltc4s* specifically ablates the ability of tuft cells to generate CysLTs but does not alter their numbers, transcriptional profile, or potential to sense environmental danger signals and respond with generation of eicosanoids.

### *Alternaria*-induced ILC2 and DC expansion depends on tuft cell-derived CysLTs

We have previously shown that tuft cells are directly activated by aeroallergens to generate CysLTs (*16*). Here we confirm the physiological importance of this finding by showing that tuft cell-specific deletion of *Ltc4s* leads to significant decrease of overall CysLT generation in the naïve nasal mucosa (**Fig. 4F**). To determine how tuft cell-derived CysLTs regulate the airway immune response to inhaled allergens, we administered the aeroallergen *Alternaria* to tuft cell-specific CysLT-ablated *Chat^Cre^Ltc4s^fl/fl^* mice and *Ltc4s^fl/fl^* controls (**Fig. 5A**). Overall *Alternaria-*induced lung inflammation was reduced in *Chat^Cre^Ltc4s^fl/fl^* mice (**Fig. 5B**), confirming the importance of tuft cell-derived CysLTs in this response.

**Fig. 5.**
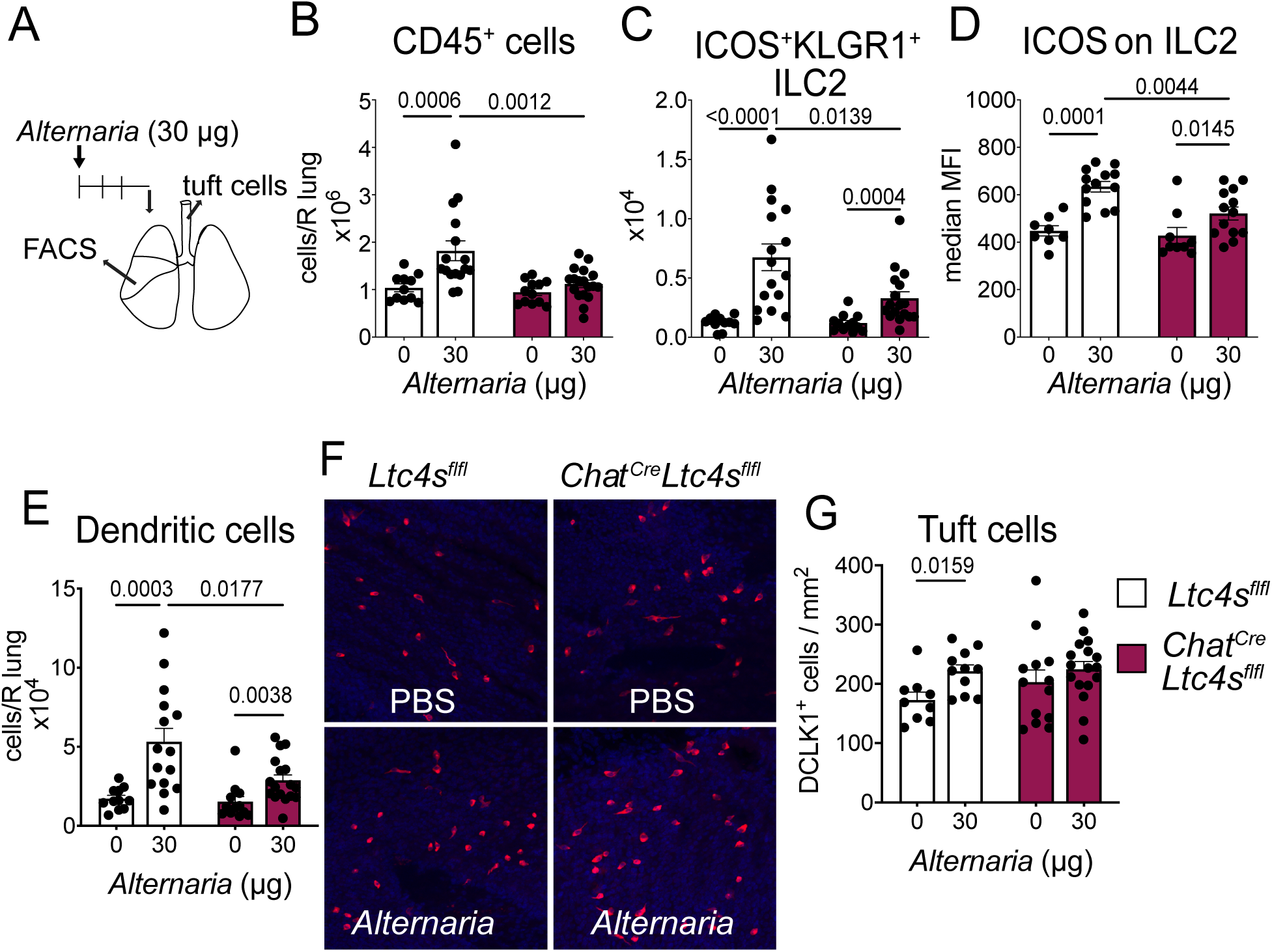
Tuft cell-derived CysLTs regulate ILC2 expansion in the lung. (**A-B**) *Ltc4s^fl/fl^* and *Chat^Cre^ Ltc4s^fl/fl^* mice were given a single intranasal administration of *Alternaria.* Tracheal tuft cells and lung inflammation were evaluated 72 hours after challenge (**A**). Flow cytometry analysis of dispersed lung cells showing total hematopoietic cell numbers (CD45^+^) (**B**), ILC2 numbers (CD45^+^lin^−^Thy1.2^+^KLRG1^+^ICOS^+^) (**C**), KLRG1 MFI (**D**) and DCs (CD45^+^SiglecF^-^CD11c^+^MHCII^+^) (**E**). Tracheal tuft cells were identified in whole tracheal mounts by DCLK1 immunoreactivity. Representative images (**F**) and quantitation of tuft cell numbers in the trachea (**G**). Data are means ± SEM, from at least three independent experiments, each dot represents a separate mouse, Mann Whitney U-test p values < 0.05 are indicated.

Our exogenous IL-25+LTC_4_ inhalation data suggested that ILC2s and DCs were prominently affected by CysLTs. Consistent with that finding, both local expansion of KLRG1^+^ICOS^+^ ILC2s and their expression of the activation marker ICOS were reduced in *Alternaria-*challenged *Chat^Cre^Ltc4s^fl/fl^* mice (**Fig. 5C, D**). Similar to our pharmacologic model, we found a reduction in the recruitment of DCs to the lung (**Fig. 5E**). Thus, in an aeroallergen-triggered airway inflammation model where CysLTs are produced early in the tissue response, the targeted deletion of CysLT generation in tuft cells diminishes the local activation and expansion of two critical innate immune cell types. *Alternaria-*induced lung inflammation is also reduced in the absence of IL-25 signaling (*20*) at the same time point. Thus, to account for the IL-25 component of the LTC_4_/IL-25 synergy and determine whether the LTC_4_/IL-25 synergy requires an intact LTC_4_ biosynthetic pathway in tuft cells, we administered LTC_4_ and IL-25 alone or in combination (as in **Fig. 1A**) to *Chat^Cre^Ltc4s^flfl^* mice. Lung eosinophilia, ICOS^+^KLRG1^+^ ILC2 expansion and DC expansion were induced in the *Chat^Cre^Ltc4s^flfl^* mice in a synergistic fashion similar to what we observed in WT mice (**Fig. S6**). To further probe if IL-25 signaling might be disrupted in the absence of endogenous CysLTs and account for all possible sources of LTC_4_, we administered a higher dose of IL-25, sufficient to induce *in vivo* activation of ILC2s (*9*) and DCs (*42*), to WT and mice with germline deletion of *Ltc4s*. Intranasal IL-25 (500 ng) induced ILC2 expansion and activation as well as DC recruitment and tuft cell number increase in the trachea, all of which were intact in *Ltc4s^-/-^* mice (**Fig. S7**). The preserved response to IL-25 in the absence of LTC_4_S suggests that IL-25 autocrine signaling in tuft cells is not a major contributor to their proinflammatory function via CysLT production. This suggests that IL-25 and CysLTs are generated by tuft cells in response to aeroallergen activation and synergistically activate innate immune cells to promote lung inflammation.

*Alternaria* inhalation leads to both lung inflammation and tracheal epithelial remodeling with tuft cell expansion that is dependent on endogenous CysLTs (*20*). Here, we found that *Alternaria* inhalation led to expansion of tuft cells in the trachea of WT *Ltc4s^fl/fl^* mice while no tuft cell expansion was observed in *Chat^Cre^Ltc4s^fl/fl^* mice (**Fig. 5F, G**). Tuft cell-specific deletion of CysLTs was associated with a higher baseline number of tuft cells, which is consistent with our previous data in *Ltc4s^-/-^* mice and points to the tuft cell as the major source of CysLTs that regulate the tracheal remodeling. Collectively, we demonstrate that endogenously generated CysLTs from tuft cells are required for both inflammation and airway tissue remodeling in response to an inhaled aeroallergen.

### Early lung eosinophilia and systemic immune response triggered by *Alternaria* depend on tuft cell-derived CysLTs

To further define the steps of the local and systemic immune response that depend on tuft cell-derived CysLTs, we evaluated the early response to *Alternaria* 24 hours after a single intranasal inhalation (*6, 16, 43*). Bronchoalveolar lavage (BAL) eosinophilia was induced in *Ltc4s^fl/fl^* mice at 24 hours and significantly reduced in *Chat^Cre^Ltc4s^fl/fl^* mice. The low-level recruitment of neutrophils at this time point was not affected by the deletion of *Ltc4s* in tuft cells (**Fig. 6A**). The 70% reduction of BAL eosinophilia in *Chat^Cre^Ltc4s^fl/fl^* mice was similar to the reduction we had previously reported in tuft cell-deficient *Pou2f3 ^-/-^* mice (*16*). Mice with germline deletion of *Ltc4s* had a similar degree of reduction in eosinophilia as *Chat^Cre^Ltc4s^fl/fl^* mice, with marginal effect on the variable rapid lung neutrophil recruitment (**Fig. 6B**). Since *Alternaria* inhalation is also associated with resident innate mast cell activation, we also evaluated mast cell-deficient mice (*43*). Mice with Mcpt5Cre-mediated diphtheria toxin ablation of innate mast cells showed a degree of reduction in BAL eosinophilia that was similar to mice with tuft cell-specific and germline deletion of *Ltc4s* (**Fig. 6B**).

**Fig. 6.**
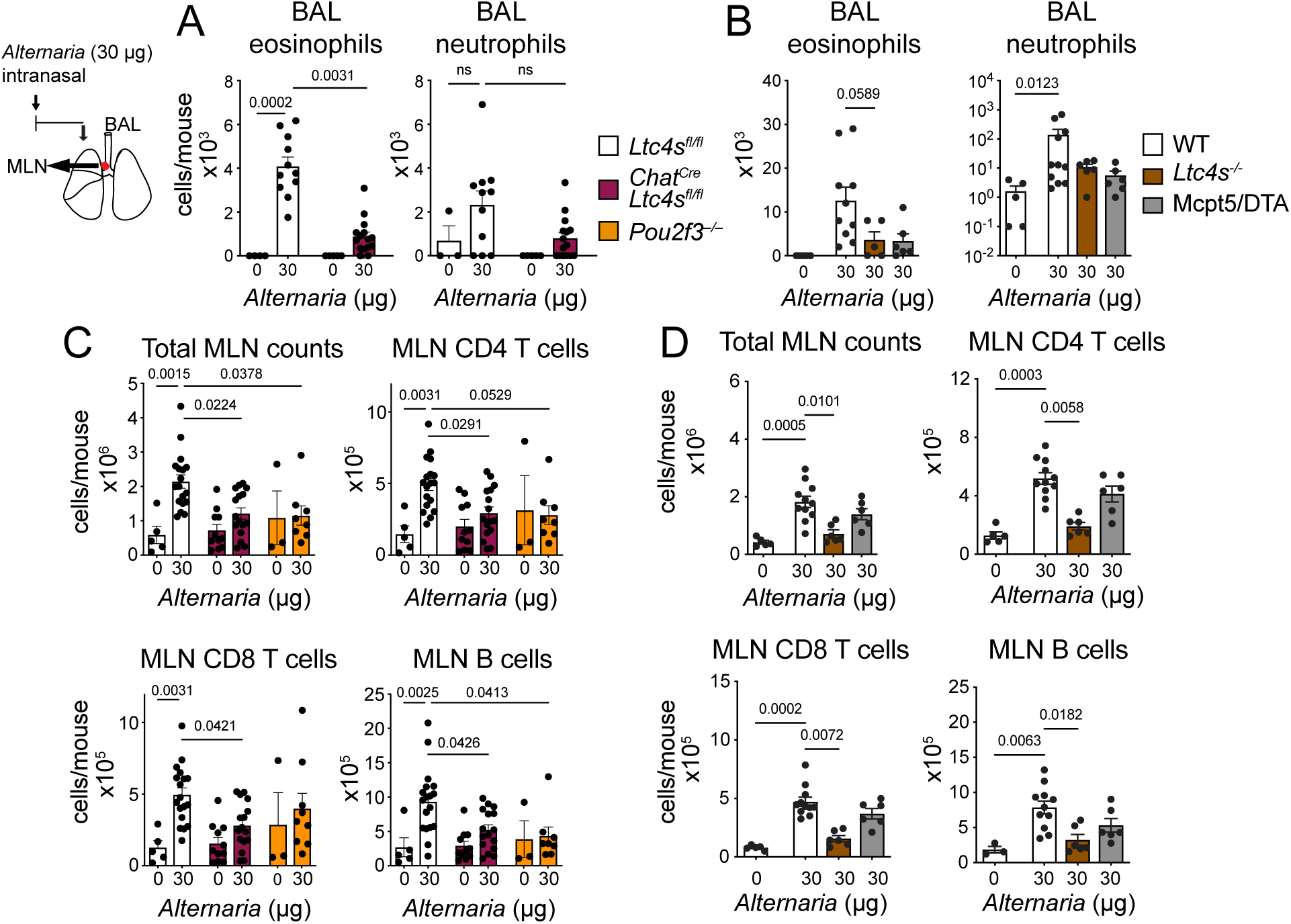
Tuft cell-derived CysLTs contribute to *Alternaria*-induced early eosinophilia and systemic immunity. WT (C57BL/6 and *Ltc4s^fl/fl^)*, *Chat^Cre^Ltc4s^fl/fl^, Pou2f3^−/−^* (**A, C**), *Ltc4s^−/−^* and Mcpt5/DTA mice (**B, D**) were given a single intranasal dose of *Alternaria* and were evaluated 24 hours after the challenge. The total number of cells in the BAL lavage were counted, and differential subsets were evaluated using Diff-Quick (**A, B**). Lung draining mediastinal lymph nodes (MLN) were harvested and the LN cell subsets were evaluated by FACS (**C, D**). Data are means + SEM pooled from ≥ 2 independent experiments, each dot is a separate mouse, p values from Kruskal Wallis test with Dunn’s multiple comparison correction.

To evaluate the relative contribution of tuft cell-derived CysLTs to the systemic immune response, we assessed the lymph node response 24 hours after a single dose of *Alternaria.* We compared mice with tuft cell-specific deletion of *Ltc4s* to tuft cell-deficient *Pou2f3 ^-/-^* mice to account for differences attributable to other tuft cell-derived mediators, including IL-25 and PGD_2_. The *Alternaria-*elicited lung draining lymph node hyperplasia was significantly reduced in *Chat^Cre^Ltc4s^fl/fl^* and *Pou2f3 ^-/-^* mice 56% and 53% respectively, pointing to a role of tuft cell-derived CysLTs in systemic immunity. We found no significant difference between tuft cell- deficient mice and mice with tuft cell-specific deletion of *Ltc4s* (**Fig. 6C**). Disrupted CysLT generation in tuft cells had a generalized effect on the proliferation and recruitment of CD4^+^ and CD8^+^ T cells and B cells (**Fig. 6C**), which mirrored the reduction in the lymph node response found in *Ltc4s^-/-^* mice with germline deletion of *Ltc4s* (**Fig. 6D**). Mast cell-deficient mice had a borderline but non-significant reduction in lymph node hyperplasia, suggesting that the tuft cell- generated LTC_4_ accounts for the CysLT-dependent effects in the lymph nodes.

In summary, we find that despite the low densities of tracheal tuft cells relative to the nasal mucosa, tuft cell-derived CysLTs are required for eosinophilic inflammation throughout the lungs and contribute to the development of the systemic immune response to aeroallergens.

## DISCUSSION

Tuft cells are sentinel airway epithelial cells positioned to sense inhaled antigens with apical tuft-like projections extending into airway lumen (*44*). The functions of these unique cells were obscure until they were firmly established as initiators of type 2 immunity in the intestine. Our results implicate them as important drivers of aeroallergen-initiated type 2 inflammation in the airways. Tuft cells are the dominant source of IL-25 in the airways, and IL-25 signaling is required for the development of type 2 inflammation in the lung after *Alternaria* inhalation (*20, 21*). Here, we demonstrate that the tuft cell-derived mediators LTC_4_ and IL-25 synergize to promote type 2 inflammation in the lung. Although signaling through CysLT_1_R drives tissue eosinophilia, signaling through both CysLT_1_R and CysLT_2_R – the latter resistant to available pharmacotherapy – is required for the full synergistic effect of LTC_4_ and IL-25 on type 2 lung inflammation. Finally, CysLT production by tuft cells in response to the aeroallergen *Alternaria* drives early eosinophilia, lymph node hyperplasia, and innate immune cell activation. These findings uncover how inflammatory lipids, previously not known to be produced by the epithelium, synergize with epithelial ’alarmin’ cytokines to drive cascades of type 2 inflammation in response to inhaled aeroallergens.

Tuft cells in the upper and lower airways have gained recognition as sensors of bitter tasting agonists and bacterial compounds (*45–47*), formylated bacterial peptides (*48*) and ATP (*16, 45, 49, 50*). ATP signaling through the P2Y2 receptor in tuft cells is engaged by the aeroallergen *Alternaria*, leading to generation of CysLTs (*16*). Consistent with that mechanism, tuft cell-deficient *Pou2f3^-/-^* mice have reduced *Alternaria-*triggered CysLT generation and reduced airway eosinophilia (*16*). Here, we used a mouse with a tuft cell-targeted deletion of *Ltc4s,* the terminal CysLT biosynthetic enzyme, to ablate the CysLT pathway specifically in these cells. Deletion of *Ltc4s* did not alter the homeostatic transcriptional profile of airway tuft cells or their ability to respond to ATP and allergen activation with generation of the lipid mediator PGD_2_, and did not alter their capacity to generate IL-25. Tuft cell-specific deletion of CysLT production impaired both early *Alternaria-*induced eosinophilia in the lung and the systemic response to this aeroallergen. Surprisingly, the reduction in airway inflammation was not further reduced in tuft cell deficient *Pou2f3^-/-^* mice (which lack tuft cell-derived CysLTs and IL-25) compared to *Chat^Cre^Ltc4s^fl/fl^* mice. This appeared inconsistent with our previous findings that IL-25 signaling is required for the full development of lung inflammation triggered by *Alternaria* (*20*). We considered that this might be due to direct activation of tuft cells by IL-25 through IL17RB for CysLT generation. However, high dose IL-25 induced lung inflammation, and the synergy of low dose IL-25 and LTC_4_ were not diminished in mice with disrupted CysLT generation. This suggests that LTC_4_ and IL-25 are likely generated concurrently when tuft cells are activated by allergens. The level of IL-25 in this model is likely too low to detect in biological fluids but sufficient to synergize with the high levels of CysLT generated in response to *Alternaria* (*6, 51*). Thus, in this *Alternaria*-initiated airway inflammation model, IL-25 signaling requires the presence of CysLTs to trigger ILC2 activation, lung eosinophil recruitment, and DC activation and migration to the lymph nodes. This system of lipid mediator synergy with IL-25 can be engaged to promote type 2 inflammation and could be engaged when the epithelial cytokine IL-25 is produced at low levels.

Consistent with the hypothesis that CysLTs might be required for optimal IL-25 signaling in allergic inflammation, we found that LTC_4_ can render low dose non-pathogenic IL-25 into a strong inducer of type 2 airway immune responses. Low-dose IL-25 was insufficient to induce lung inflammation but synergized with LTC_4_ to activate ILC2s and DCs and to potently induce type 2 eosinophilic lung inflammation. Neuromedin U similarly synergizes with IL-25 to greatly potentiate type 2 inflammation (*9*). These studies demonstrate that several mediators derived from tissue resident cells promote the effect of IL-25 on type 2 immunity and suggest that IL-25 effects are tightly regulated, likely to limit their pathogenetic potential. IL-25 and LTC_4_ have an additive effect on intestinal ILC2 activation *in vitro* (*32*) but their effect *in vivo* far exceeds the effect of either mediator alone. Beyond ILC2s, several immune cell subsets were activated and increased in numbers, including DCs, with a specific increase in CD301b^+^ DCs and CD4 T cells. Finally, IL-4, IL-5, IL-13 and IL-6 were potently induced by IL-25 + LTC_4_, with moderate induction of TNF-α and no effect on interferon γ or IL-17 type cytokines. Thus, this lipid mediator/epithelial cytokine synergy induces activation of several immune cell types and potently skews tissue inflammation towards type 2 immune response.

Our findings of receptor requirements for both CysLT_1_R and CysLT_2_R signaling for the development of type 2 inflammation in response to LTC_4_+IL25 highlight the complexity of the CysLT signaling system. CysLT_1_R and CysLT_2_R are broadly expressed in hemopoietic cells of the innate (macrophages, monocytes, eosinophils, basophils, mast cells, DCs) and adaptive (T cells, B cells) immune system, suggesting multiple sites of possible cooperative functions in inflammation (*52*). Among the innate immune cells specifically activated by IL-25 + LTC_4_, ILC2s and DCs express both CysLT_1_R and CysLT_2_R (*5, 7, 53, 54*), and both cell types also express IL17RB (*9, 34*). Despite the reported broad expression of both CysLT receptors, here we find that CysLT_1_R is the dominant receptor mediating lung eosinophilia and DC recruitment in response to LTC_4_ + IL-25. This is consistent with the major effect of CysLT_1_R in mediating the synergy of IL-33 and LTC_4_ on lung eosinophilia (*7*) and lung ILC2 activation (*5, 7*). While the potentiation effect of LTC_4_ on IL-33-induced ILC2 activation and lung eosinophilia is independent of CysLT_2_R signaling, we find that both CysLT_1_R and CysLT_2_R are required for induction of type 2 cytokines in the lung. This is consistent with the effect of LTC_4_ on eosinophilic lung inflammation in mice sensitized with ovalbumin. In this model, LTC_4_ induces ILC2 expansion and type 2 cytokines IL-5 and IL-13, an effect dependent on both CysLT_1_R and CyLT_2_R (*30, 55*). Thus, the effect of LTC_4_ likely depends on the specific cellular milieu in the vicinity of the LTC_4_ source. Interestingly, we find that both CysLT_1_R and CysLT_2_R are required for the expansion of the inflammatory ILC2 subset of KLRG1^+^ICOS^+^ ILC2s. This specific subset of “inflammatory” ILC2s was previously described as specifically induced by high dose IL-25 and requires signaling through IL17RB (*9, 34*). Here, we observe that LTC_4_ facilitates the induction of these inflammatory ILC2s. The fact that CysLT receptor deletion sharply diminishes their numbers in the lung is consistent with their dependence on IL17RB signaling, and with the exquisite requirements for physiologically generated IL-25 to synergize with CysLTs. Thus, the *in vivo* selectivity of LTC_4_ might differ depending on the tissue distribution of the cellular source, the epithelial cytokines generated in concert with these lipid mediators, and the frequency of CysLT receptor expressing cells and their proximity to the ligand source (*53, 56–58*).

*Alternaria* and house dust mite inhalation are associated with an increase in the frequency of tracheal tuft cells dependent on endogenous CysLTs and only partially dependent on *Stat6* signaling (*20*). Here we find that mice with tuft cell deletion of *Ltc4s* phenocopy the germline *Ltc4s^-/-^* mice for the absence of tuft cell expansion, suggesting that these cells are a major contributor to the endogenous CysLTs needed for this function. Previously, we had shown that exogenous LTE_4_, the most stable CysLT, increases the frequency of tracheal tuft cells in a *Stat6-* independent manner mediated by *Cysltr3* (*20*). suggesting a CysLT-dependent mechanism of epithelial remodeling independent of classical IL-13-driven mechanisms. Now, we find that tuft cells in the trachea are not induced in our pharmacologic model despite strong induction of lung IL-13 and concomitant goblet cell metaplasia in the lung. This supports the observation that tracheal tuft cell expansion in the airways is not directly dependent on IL-13, despite the dominant role of this cytokine in goblet cell metaplasia in the airways and in intestinal tuft cell expansion. The dichotomy of IL-13-dependent goblet cell metaplasia and IL-13-independent tracheal tuft cell expansion – even in the presence of LTC_4_ – further highlights the unique role of each member of the CysLT family, where LTC_4_ acts in concert with epithelial cytokines to induce robust type 2 inflammation, while LTE_4_ has an independent effect on tracheal remodeling for tuft cell expansion (*20*).

While the pharmacological synergy of IL-25 with LTC_4_ for type 2 immunity suggest that both are central mediators produced by tuft cells, we cannot exclude the possibility that LTC_4_ from tuft cells might have broader effects on immune responses beyond this specific interaction. Tuft cells are also prominent sources of acetylcholine, which can be generated in response to formylated peptides (*48*). Recent studies suggest that a population of *Chat*-expressing ILC2 that also expresses cholinergic receptors is specifically induced by *Alternaria* in the lung (*39*). Thus, an interaction between acetylcholine and CysLTs might further augment the activation of ILC2s. Interestingly, although both *Chat* and *Ltc4s* are widely expressed in diverse immune subsets, we find that only tuft cells uniquely co-express both markers, further expanding the possibility of a functional importance of this co-localization.

In summary, our data provide evidence of a system of allergen-triggered responses in the airways. This system is initiated by activation of specialized airway tuft cells, which then generate potent lipid mediators that trigger the innate phase of type 2 inflammation via downstream activation of innate immune cells. Two mediators generated by tuft cells – LTC_4_ and IL-25 – cooperate for an enhancement of the inflammatory response. Notably, in mouse models, the most dramatic increase in airway tuft cell numbers is found in the recovery phase of influenza infection (*59*). In human airway diseases, CysLTs are found in the BAL of infants with RSV bronchiolitis (*60–62*) and IL-25 is induced by viral exacerbations of asthma (*63*). Beyond allergen recognition, this system may also play decisive roles in other airway diseases associated with IL-25 and CysLT production.

## MATERIALS AND METHODS

### Study design

The aims of this study were to define the role of two of the major tuft cell-derived mediators – LTC_4_ and IL-25 in driving type 2 inflammation in the airways. We used a pharmacologic model to evaluate the lung inflammation induced by LTC_4_ or IL-25 alone or in combination. Using mice with receptor-specific deletion for each CysLT receptor, we identified the two receptors responsible for mediating this response. We then developed and characterized transcriptionally a new mouse with tuft cell-specific deletion of LTC_4_ generation – *Chat^Cre^Ltc4s^fl/fl^.* We then used this mouse to specifically characterize the dependence on tuft cell-derived CysLTs in a mouse model of allergen-induced inflammation.

### Experimental Models and Mice

ChAT^BAC^-eGFP [B6.Cg-Tg(RP23–268 L19-EGFP)2Mik/J], *ChAT-IRES-Cre::frt-neo-frt* [*Chat^tm2(cre)Lowl^*], C57BL/6 and C57BL/6N-Cysltr1tm1^Ykn^/J mice were purchased from Jackson Laboratories *(Bar Harbor, ME). Ltc4s^−/−^* mice were generated on a 129Sv background and backcrossed for 15 generations onto the C57BL/6 background (*64*). CysLT_2_- (*Cysltr2^-/-^*) and CysLT_3_-receptor gene-knockout mice (*Oxgr1^-/-^* or *Cysltr3^-/-^*) were generated on a C57BL/6 and as previously described (*43, 65*). Mcpt5/DTA mice were generated by crossing mice with mast cell-specific expression of Cre recombinase (*66*) on a C57BL/6 background with ROSA-diphtheria toxin-α mice (B6.129P2-Gt(ROSA)26Sortm1(DTA)Lky/J from Jackson Laboratories). *Pou2f3^−/−^* mice were generated as described previously (*67*) and compared to C57BL/6 mice bred in-house (originally from Charles River Laboratories, Wilmington, MA). *Ltc4s^fl/fl^* mice were provided by Dr. J. Boyce and generated by Ingenious Lab. B6. *Ltc4s^fl/fl^* mice with loxP sites flanking the 1st exon of *Ltc4s* were generated by homologous gene targeting in C57BL/6 embryonic stem cells. Targeted iTL BF1 (C57BL/6 FLP) embryonic stem cells were microinjected into Balb/c blastocysts. Resulting chimeras with a high percentage black coat color were mated to C57BL/6 WT mice to generate Germline Neo Deleted mice. Homozygous *Ltc4s^fl/fl^* mice were then crossed with *Chat^Cre^* mice to generate double heterozygous mice (*Chat^Cre+^Ltc4s^fl^*), which were further mated with *Ltc4s ^fl/fl^* mice to generate two genotypes of mice: *Ltc4s^fl/fl^* and *Chat^Cre+^Ltc4s^fl/fl^* and used as littermate controls of each other.

All animals were maintained in a specific pathogen-free facility at the Brigham and Women’s Hospital; litter was weaned between 19-28 days old. All experiments were performed during the day. Pooled results include both male and female mice. The use of mice for these studies was in accordance with review and approval by the Animal Care and Use Committee of Brigham and Women’s Hospital. Mice were randomly assigned to treatment groups after matching for sex, age and genotype.

### Aeroallergen, IL-25 and LTC_4_ Challenge Protocols

Mice were anesthetized with an intraperitoneal injection of ketamine (10 mg/kg) and xylazine (20 mg/kg) for full sedation and received a single intranasal (i.n.) application of pharmacologic agonists or allergen dissolved in 20 μl PBS. For low-dose IL-25 and LTC_4_ + IL- 25 synergy experiments, mice were treated intranasally with either 1.6 mmol of LTC_4_, or 100 ng IL-25, or combination of 100 ng IL25 and 1.6 mmol of LTC_4_. LTC_4_ was stored in ethanol; the ethanol was evaporated and LTC_4_ was reconstituted in PBS for immediate application. In another set of experiments, mice were given 500 ng IL-25 intranasally. LTC_4_, IL-25 and the combination of LTC_4_ and IL-25 were administered in 20 μl volume daily for three consecutive days. Tissues were harvested 48 hours after the last intranasal administration.

For allergen challenge experiments, mice were give a single intranasal inhalation of 30 μg of *Alternaria alternata* culture filtrate. Mice were euthanized with ketamine overdose at 1 hour for nasal lavage, at 24 hours for BAL or mediastinal lymph node and at 72 hours for lung and tracheal harvesting.

### Nasal and BAL lavage

For nasal lavage, mice were euthanized 1 hour after the i.n. challenge, and the lavages were obtained through a small incision in the trachea. The nasal lavage was performed with 200 μl of PBS and the fluid proteins were precipitated with ice cold acetone. After high-speed centrifugation, the supernatant was used for CysLT and PGD_2_ measurements.

BAL for assessment of inflammation was collected by repetitive (n=3) injection and aspiration of the lungs with 0.75 ml of PBS with 1 mM EDTA through a tracheal opening. The BAL fluid was centrifuged at 500*g* for 5 min and the supernatant was discarded. Cell pellets were resuspended in 0.2 ml of PBS, and the total cells were counted manually under a microscope with a hemocytometer. For the differential cell counts of macrophages, neutrophils, and eosinophils, the BAL cells were cytospun onto fresh labeled glass slides at 500 rpm for 5 min. Air-dried slides were further stained using Diff-Quick stain kit. Cell types in a total of 200 cells were identified by morphologic criteria (*53*) and differential cell counts were performed under microscope.

### Single cell suspensions preparation

For single-cell preparations of the nasal mucosa, each euthanized mouse was decapitated, and the snout was released of surrounding bones and tissues. The nasal mucosa was exposed dorsally by cutting nasal bone located on the calvarium and ventrally by incising along the palate.

In a set of experiments with ChAT-eGFP mice used for RNAseq, the olfactory mucosa overlying the ethmoturbinates, dorsal nasal meatus and caudal half of the septum were separated from respiratory mucosa, comprised of nasal conchae and the ventral half of the septum, containing the vomeronasal organ. For all other genotypes, the whole snout, including the olfactory and respiratory areas, was processed together.

The nasal mucosa was incubated in pre-warmed PBS solution with dispase (16 U/ml) and deoxyribonuclease I (DNase I; 20 μg/ml) for 30 to 40 min at room temperature on a rotating shaker at 220 rpm. Dispase activity was reduced with Dulbecco’s modified Eagle’s medium (DMEM) and 5% fetal bovine serum (FBS). The separated nasal mucosa was incubated in Tyrode’s Solution with bicarbonate, HEPES, 0.25% BSA, without Ca^2+^, with added 48 U/mg of papain, 25 mg/mL L-cysteine and 20 μg/mL DNAse I for 40 min at 37 °C on a rotating shaker (*68*). Papain digestion was terminated with Tyrode’s Solution with HEPES and Ca^2+^ and with 2 μl/mL leupeptin (5 mg/ml). The digested tissue was vortexed for 20-30 sec, triturated thoroughly with an 18-gauge needle followed by a 21-gauge needle, passed through a 100 μm filter, and washed with FACS buffer (based on Hank’s Balanced Salt Solution (HBSS) without Ca^2+^, with added 2% FBS and 2 mM EDTA).

To isolate nasal hematopoietic cells for RNAseq, the snout was harvested as described above. The freshly isolated nasal mucosa was enzymatically dissociated with a PBS-based solution containing collagenase (650 U/ml), dispase (16 U/ml), and DNase I for 30 mins at 37°C on a rotating shaker. The digestion was terminated by addition of 1 ml of cold DMEM with 5% FBS. Digested mucosal samples were further vortexed, triturated and processed similarly to epithelial cells described previously.

For evaluation of lung inflammation, lungs were flushed with 10 ml of cold HBSS until they appeared white in color. The right lower lobe was removed, physically dissociated with a gentle MACS Dissociator in 10 ml of RPMI 1640 containing 10% FBS, followed by digestion with collagenase (500 U/ml), dispase (6.6 U/ml) and DNase I (20 μg/mL) in RPMI 1640 with 5% FCS at 37°C for 30 min with agitation at 200 rpm. The digestion was terminated by adding 10 ml of cold RPMI 1640 with 10% FBS. Cell suspensions were washed with FACS buffer and prepared for analysis by flow cytometry.

To isolate immune cells, harvested lung draining mediastinal lymph nodes were digested in PBS solution containing collagenase (100 U/ml) and DNase I (20 μg/ml) for 30 mins at 37 °C. The cell suspension was further triturated with a pipette tip, filtered, washed and further stained for flow cytometry.

### Flow cytometry and sorting

Before staining the cells, the nonspecific monoclonal antibody uptake was blocked with CD16/32 for at least 30 min. All samples were FSC-A/SSC-A gated to exclude debris, SSC-H/SSC-W and FSC-H/SSC-W gated to select single cells. Propidium iodide (PI) was used as a dead cell exclusion marker for samples submitted for sorting, while Hoechst was used for the same purpose in FACS analysis. The nasal epithelial cells were stained using monoclonal antibodies against CD45 and EpCAM for 40 min, followed by dead cell exclusion before sorting. Cells from ChAT-eGFP mice were collected in two subsets: tuft cells, identified as EpCAM^high^ GFP^+^ cells (**Fig. S3A**), and other epithelial cells that were EpCAM^high^ GFP^-^ (6). The cells were additionally gated as FSC^low^SSC^low^, and EpCAM^high^GFP^+^FSC^low^SSC^low^ cells were collected from olfactory epithelium, while EpCAM^high^GFP^+^FSC^high^SSC^high^ represented tuft cells from respiratory mucosa.

Epithelial cells from *Ltc4s^fl/fl^* and Chat^Cre^*Ltc4s^fl/fl^* mice were collected based on their expression of CD45 and three populations of cells were submitted for RNAseq (**Fig. S3B**). These populations were EpCAM^high^CD45^low^FSC^low^SSC^low^, EpCAM^high^CD45^neg^FSC^low^SSC^low^, and EpCAM^int^CD45^neg^SSC^low^ epithelial cells.

DCs were defined as CD45^+^CD11b^+^SiglecF^−^CD11c^+^MHCII^+^ cells, and macrophages were identified as CD45^+^CD11b^+^SiglecF^+^CD11c^+^ cells after excluding CD19 and TCRβ^+^ cells. (**Fig. S4C**).

To identify lung ILC2s among CD45^+^ cells, lineage-positive cells were excluded using monoclonal antibodies against CD19, CD8, GR1, Fcer1, NK1.1, CD11c, CD11b, TCRβ, TCRγδ, TER119 and CD4. ILC2 cells were identified as lin- Thy1.2^+^ cells with low forward and side scatter (**Fig. S1B**). Lung DCs were defined as CD45^+^ SiglecF^−^CD11c^+^MHCII^+^ cells and further differentiated based on their expression of CD301b and CD11b. Macrophages were identified as CD45^+^ SiglecF^+^CD11b^-^ CD11c^+^ cells, and eosinophils as CD45^+^ SiglecF^+^CD11b^+^ CD11c^-^ SSC^high^ after excluding CD19 and TCRβ^+^ cells

In lymph nodes, T cells were identified as CD8^+^ and CD4^+^ and B cells were distinguished based on their expression of B220.

### Ex vivo stimulation of nasal tuft cells

Sorted EpCAM^high^CD45^low^FSC^low^SSC^low^ cells from *Chat^Cre^, Ltc4s^fl/fl^* and *ChAT^Cre^Ltc4s^fl/fl^* mice were plated in DMEM-based EpC proliferation media, which includes DMEM/F-12, 5% Nu-Serum, 100 μg/mL Penicillin-Streptomycin (Gibco) and 0.25 μg/ml Amphotericin B. Sorted CD45^+^ cells from *Ltc4s^fl/fl^* and *Chat^Cre^Ltc4s^fl/fl^* mice were reconstituted in R10 media (RPMI medium 1640 containing 10% (vol/vol) FBS, 2 mM L-glutamine, 0.1 mM nonessential amino acids, penicillin (100 units/ml), streptomycin (100 μg/ml). The cells were plated into 96-well culture plates at a concentration of 400,000 cells/ml in 200 μL of media per well for epithelial cells and 1.2x10^6^-2.3x10^6^cells/1 ml in 200 μL for CD45^+^ cells. The cells were rested overnight and stimulated with 1 mM A23187 or 0.5 mM ATPγS in 50 μl of HBSS with Ca^2+^ and Mg^2+^ for 30 min. After stimulation, the plate was centrifuged at 350 × g for 5 min at 4 °C, and the supernatants were retained for assays of CysLTs and PGD_2_.

### CysLT and PGD_2_ detection

CysLT and PGD_2_ generation in the supernatants of tuft cells, CD45^+^ sorted cells, and acetone-precipitated nasal lavage fluid were measured by commercially available enzyme ELISAs according to the manufacturer’s protocol (both kits from Cayman). The assays are based on competition between CysLTs/ PGD_2_ and CysLT-acetylcholinesterase/ PGD2-acetylcholinesterase conjugate for a limited amount of CysLT/PGD_2_ ELISA Monoclonal Antibody. The lower limit of detection for CysLT ELISA was 60 pg/mL and the following reported reactivity: Leukotriene C_4_ (100%), Leukotriene D_4_ (100%), Leukotriene E_4_ (79%), 5,6-DiHETE (3.7%), Leukotriene B_4_ (1.3%), 5(S)-HETE (0.04%), Arachidonic Acid (<0.01%). The lower limit of detection for PGD_2_ ELISA was 19.5 pg/mL and the following reported reactivity: Prostaglandin D_2_ (100%), Prostaglandin F_2α_ (92.4%), Prostaglandin J_2_ (21.6%), Prostaglandin E_2_ (2.86%), Thromboxane B_2_ (2.54%), 11β-Prostaglandin F_2α_ (1.99%), 8-iso Prostaglandin F_2α_ (1.90%), Prostaglandin A_2_ (0.72%), 12(S)-HHTrE (0.16%), 6-keto Prostaglandin F_1α_ (0.05%), (13,14-dihydro-15-keto Prostaglandin D_2_) 0.02%, Arachidonic Acid ( <0.01%), Leukotriene D_4_ (<0.01%), tetranor-PGDM (<0.01%), tetranor-PGEM (<0.01%), tetranor-PGFM <0.01%) and tetranor-PGJM (<0.01%).

### Cytokine detection in the lung

For protein extraction, the frozen lobes of the right lungs were mechanically homogenized in 300 μl of T-PER protein extraction buffer supplemented with appropriate amount of protease inhibitor cocktail (cOmplete^™^, Mini Protease Inhibitor Cocktail, Sigma Aldrich; 10 mL T-Per buffer/1 tablet). The suspensions were spun, and the supernatants were collected. The protein concentration was measured with commercially available bicinchonic acid (BCA) protein assay kit (ThermoFisher) according to manufacturer’s instructions.

The cytokine concentrations were measured with LEGENDplex T-helper Cytokine Panel (Biolegend) in V-bottom plates, following the manufacturer’s instructions. In brief, supernatants were incubated for 2 hours with fluorescence-encoded beads, differentiated by size and internal fluorescence intensities to detect 12 cytokines. The cytokine panel included IFN-γ, IL-5, TNF-α , IL-2, IL-6, IL-4, IL-10, IL-9, IL-17A, IL-17F, IL-22, IL-13 After washing, a biotinylated detection antibody cocktail was added, followed by incubation with streptavidin-phycoerythrin. Fluorescent signal intensities were then measured on 5-Laser BD FACSAria Fusion.

### Histochemistry and quantitative assessment of goblet cell numbers

For histochemical evaluation, the left lung was embedded in glycolmethacrylate. Tissue sections, 2.5 μm thick, were assessed by PAS for quantitation of mucin-containing goblet cells and by Congo red reactivity for quantitation of eosinophil recruitment. Slides were counterstained with hematoxylin for general morphologic examination. All histologic assessments were done in a blinded fashion by a single investigator.

The number of PAS-reactive cells for each animal was enumerated from 3-6 10-20x digital photographs spanning 2-13 mm basement membrane over the 4 coronal sections. This area was divided by the length of basement membrane to define the average thickness of the submucosal space.

### Confocal microscopy of whole tracheal mounts and quantitative assessment of tuft cells

The harvested tracheas were fixed with 4% PFA, washed with PBS and permeabilized in a PBS-based blocking buffer containing 0.1% Triton X-100, 0.1% saponin, 3% bovine serum albumin, and 3% normal donkey serum for at least 3 hours. The tracheas were further incubated with primary rabbit anti-DCLK1 antibody added to the permeabilizing solution at 4°C for 48 to 72 hours. The samples were then washed with PBS containing 0.1% Triton X-100 for 3 hours and transferred to PBS containing donkey anti-rabbit secondary antibody, Alexa Fluor Plus 594 and Hoechst 33342 for nuclear stain. Tracheas were longitudinally split into two halves and embedded with the epithelial surface facing upward using a glycerol-based cover slipping solution. Images were acquired at the Brigham and Women’s Hospital Confocal Microscopy Core Facility using the Zeiss LSM 800 with Airyscan confocal system on a Zeiss Axio Observer Z1 inverted microscope with 10× Zeiss [0.30 numerical aperture (NA)], 20× Zeiss (0.8 NA), and a 63× Zeiss oil (1.4 NA) objectives. The number of tuft cells for each animal was enumerated from 6 to 8 photographs corresponding to 0.6-1 mm^2^ taken from the same areas of each trachea starting at the distal end in proximity to the carina. The total number of DCLK1 immunoreactive cells (ranging from 131 to 205 tuft cells in unchallenged mice to 144 to 296 tuft cells in challenged mice) was divided by the area captured in the focal plane to define the number of DCLK1^+^ cells per square millimeter. DCLK1 immuno-reactive cells and focal plane area were evaluated using ImageJ (National Institutes of Health, Bethesda, MD). Separate images were collected for each fluorochrome and overlaid to obtain a multicolor image.

### Low input RNA sequencing

#### Preprocessing

BCL files were converted to merged, de-multiplexed FASTQ files using the Illumina Bcl2Fastq software pack-age v.2.17.1.14. Paired-end reads were mapped to the UCSC mm10 mouse transcriptome using Bowtie57 with parameters ‘-q–phred33-quals -n 1 -e 99999999 -l 25 -I 1 -X 2000 -a -m 15 -S -p 6’, which allows alignment of sequences with one mismatch. Gene expression levels were quantified as transcript-per-million (TPM) values by RSEM58 v.1.2.3 in paired-end mode. For each sample, we determined the number of genes for which at least one read was mapped, and then excluded all samples with fewer than 10,000 detected genes. Only one sample was excluded in this way. Computational pipelines for RNA seq analysis were implemented as described elsewhere (*11, 24*).

Briefly, variable genes were selected using logistic regression fit to fraction of samples in which each transcript was detected, using the log of total transcripts for each gene as a predictor. Outliers from this curve are genes that are expressed in a lower fraction of samples than would be expected given the total number of reads mapping to that gene, that is, treatment condition or state-specific genes. We used a threshold of deviance <−0.2, producing a set of 2307 variable genes. We restricted the expression matrix to this subset of variable genes, and took the variance stabilizing transformation log2(TPM +1) to generate the digital gene expression (DGE) matrix that was used for all downstream analysis. We performed dimensionality reduction on the DGE matrix using PCA. Values were centered and scaled before input to PCA, which was implemented using the R function ‘prcomp’ from the ‘stats’ package. The data were visualized in Fig. 3 using the first two principal components, which accounted for 29% and 10% of the variance respectively.

#### Comparison to known tuft cell signature

Tuft cell marker genes were derived from the ‘consensus’ signature defined with single-cell RNA sequencing (*11*). Heatmaps were generated using the ‘pheatmap’ R package.

#### Correlation of nasal epithelial groups

Pairwise Pearson correlations between all samples were calculated using the R function ‘cor’ from the ‘stats’ package (**Fig. S3D**). To identify the similarity between the putative prospectively isolated tuft cells (*Ltc4s^fl/fl^* EpCAM^high^ CD45^low^), the mean expression of each gene across the *Ltc4s^fl/fl^* EpCAM^high^ CD45^low^ tuft cell-enriched samples, and used this as a reference signature to calculate Pearson correlations with other samples. We visualized the correlation of each sample to the *Ltc4s^fl/fl^* EpCAM^high^ CD45^low^ consensus signature, grouped by sample population, and ranked these by their mean correlation.

### *Ltc4s* expression in neurons

Expression data for mouse neuronal populations was downloaded from the Mouse Brain Atlas (http://mousebrain.org/downloads.html) and visualized using the ‘ggplot2’ package in R.

### Statistics

Analysis was performed with GraphPad Prism software (version 8). Nonparametric two-sided Mann–Whitney and unpaired T tests were used to determine significance in pairwise comparison of responses in the *in vivo* and *ex vivo* stimulation models. For experiments with ≧4 group comparisons, the overall significance was determined using a one-way ANOVA and pairwise comparison was performed with Kruskall-Wallis tests with Dunn’s correction. A value of p <0.05 was considered significant. Sample sizes were not predetermined by statistical methods.

**TABLE 1.**
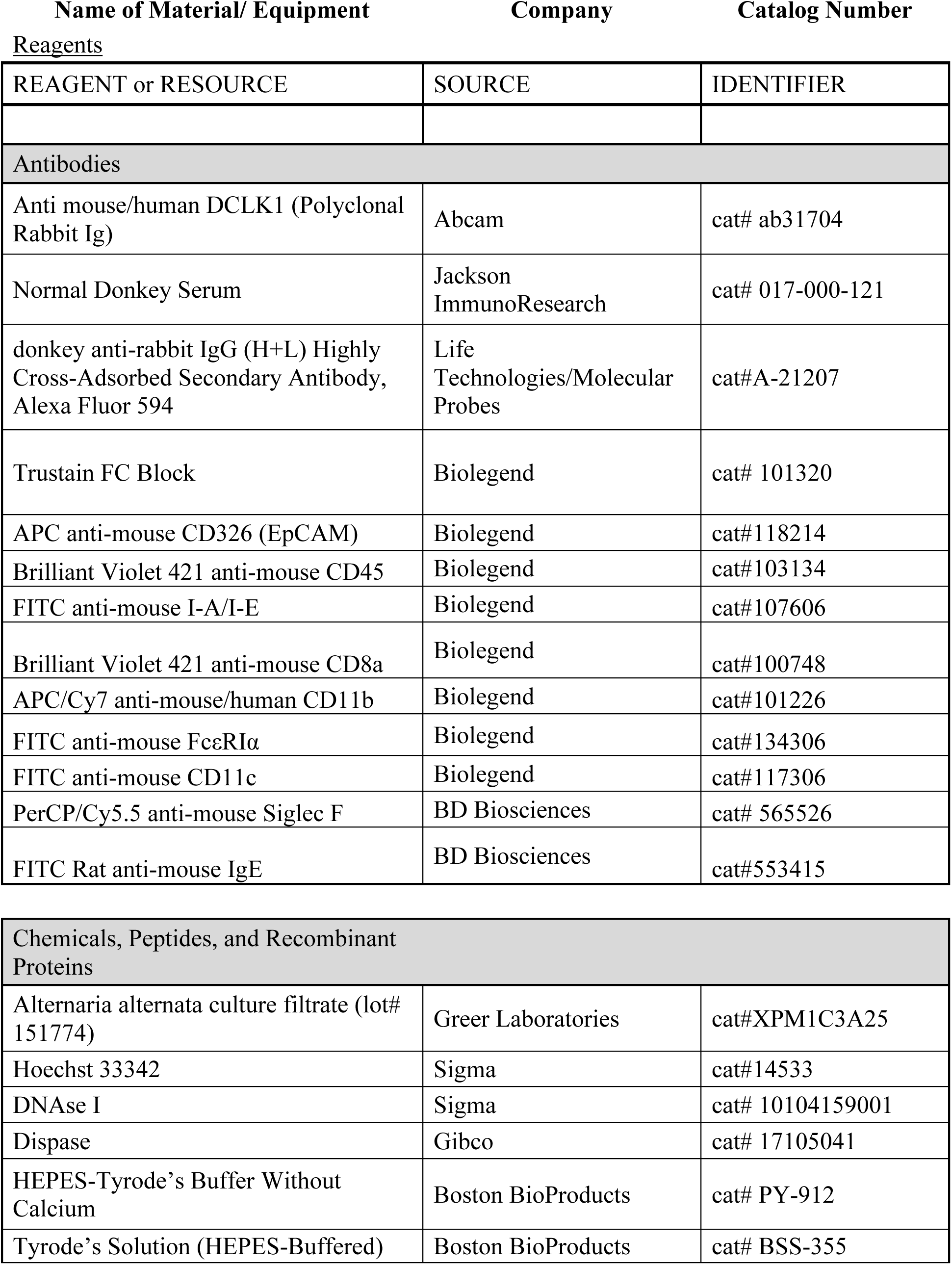
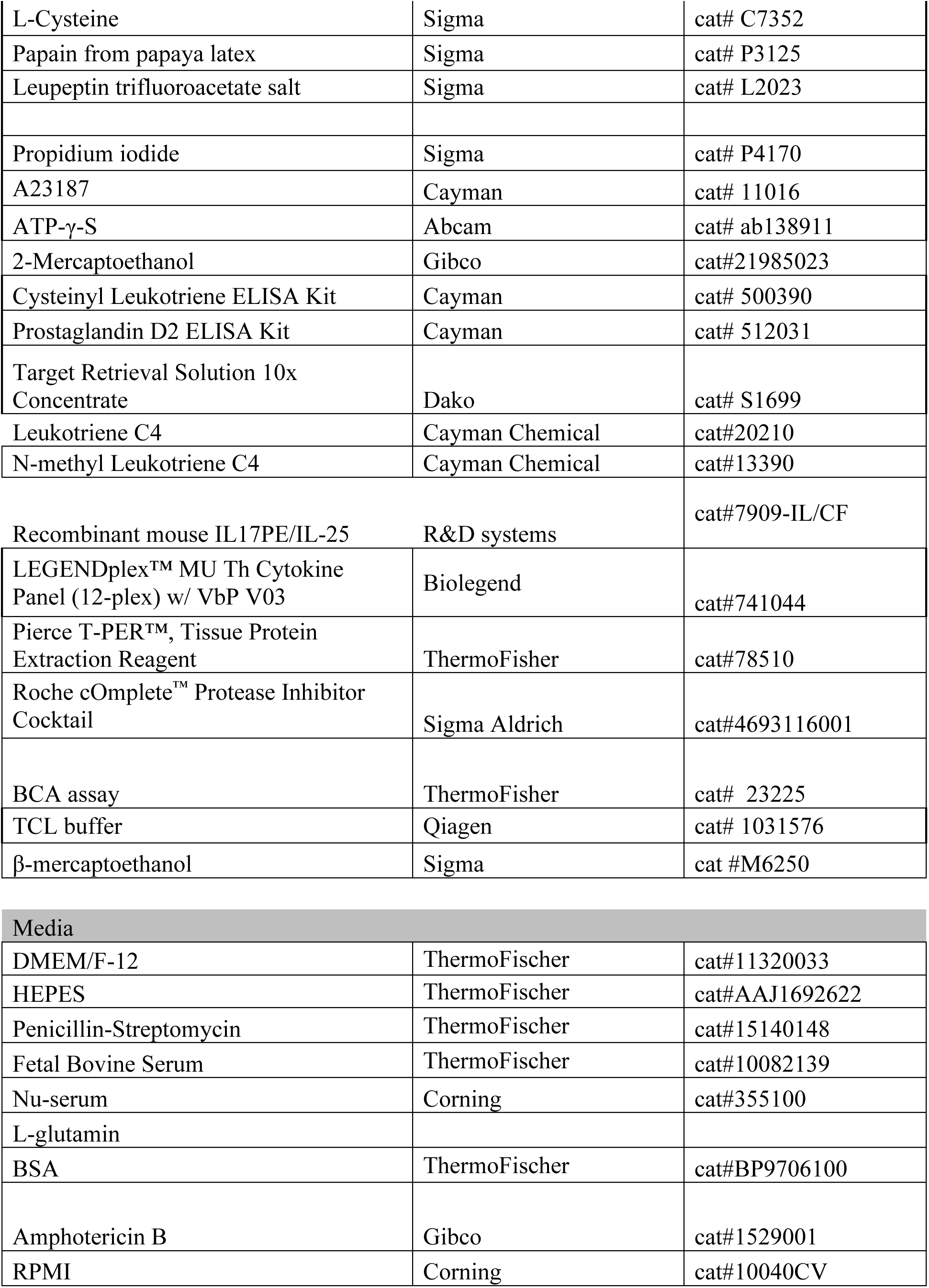
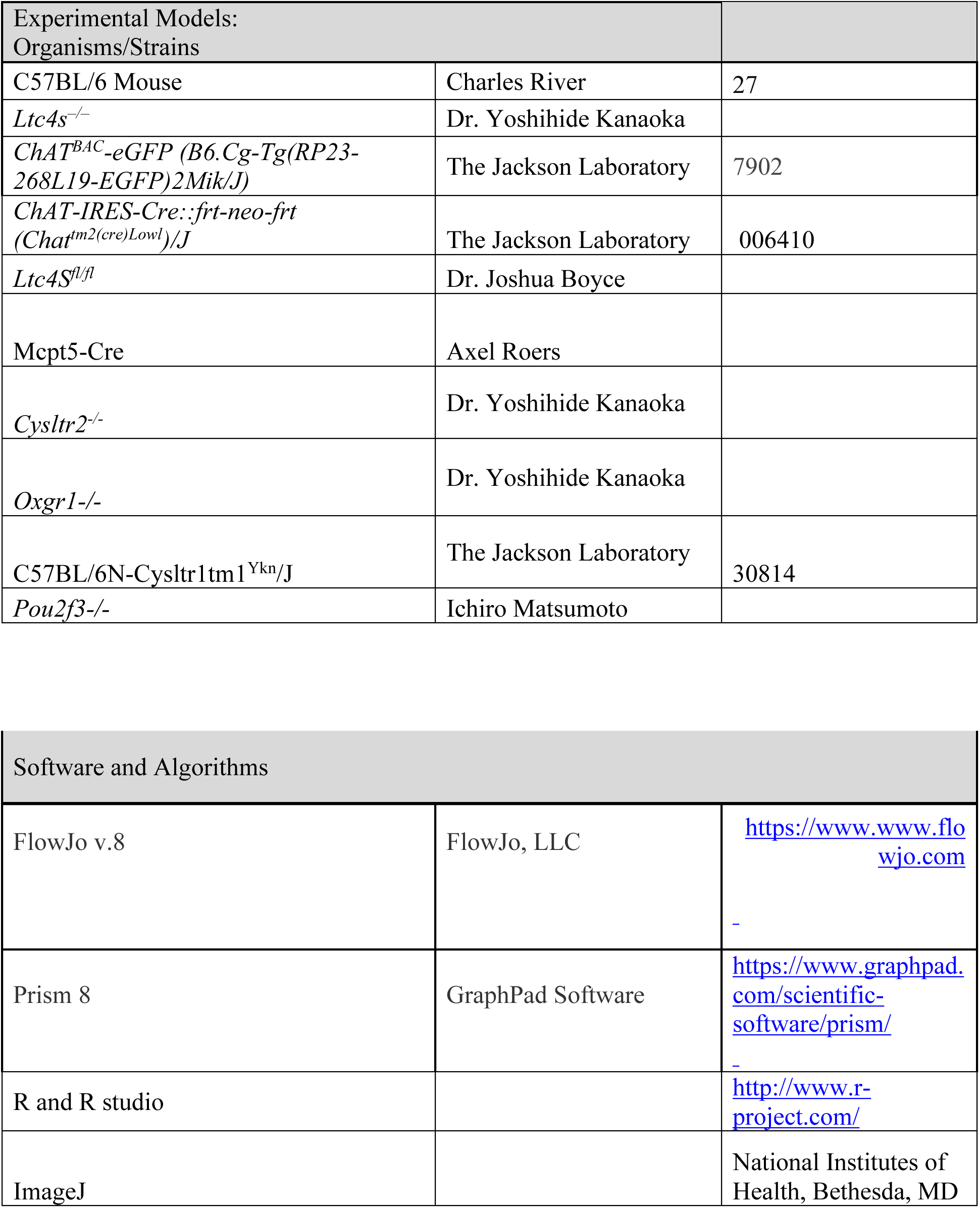

## SUPPLEMENTARY MATERIALS

**Fig. S1.**
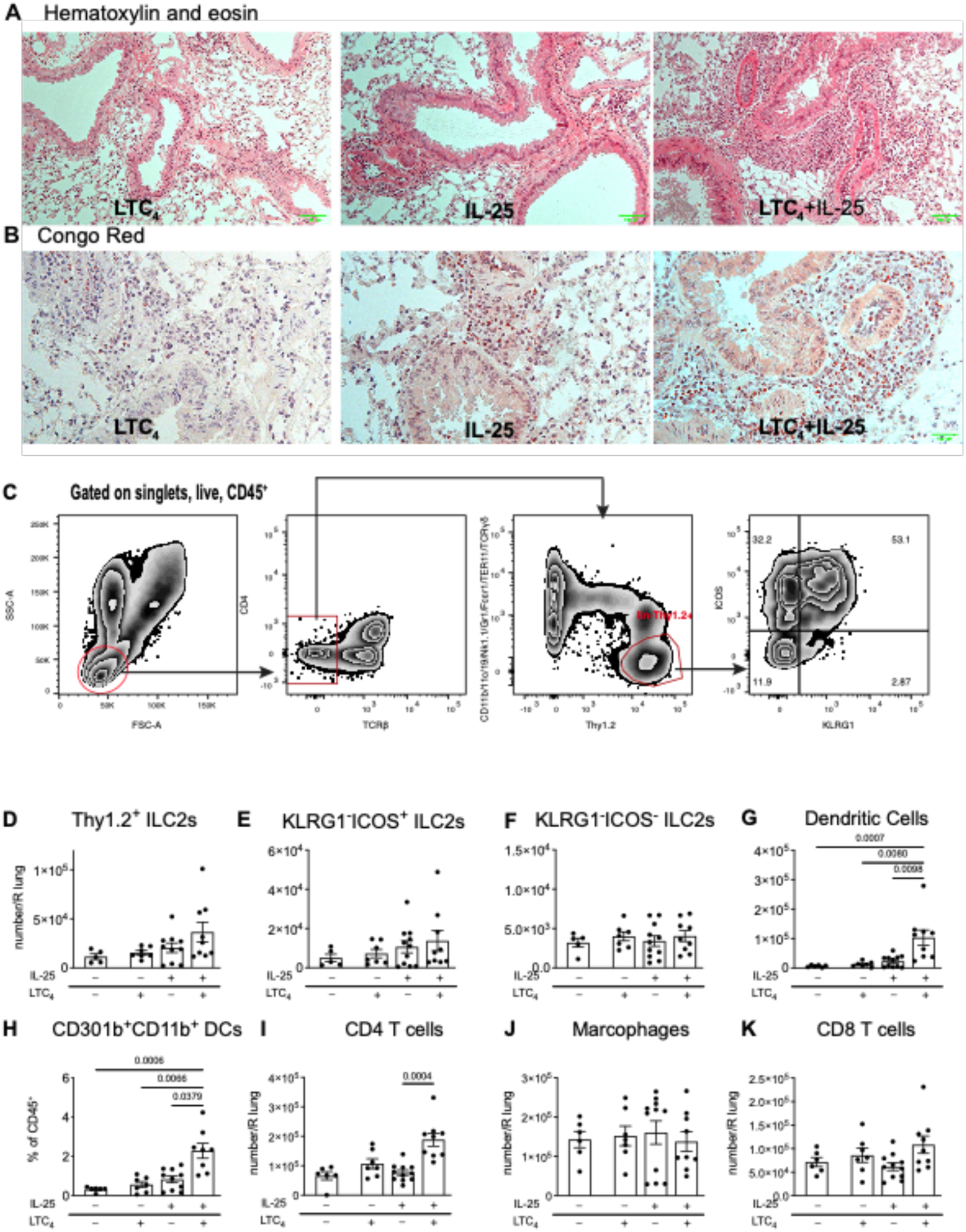
LTC_4_ and IL-25 synergize for airway type 2 lung inflammation. (A) WT (C57BL/6 and *Ltc4s^fl/fl^)* or *Chat^Cre^Ltc4s^fl^*^/fl^ mice were given three daily inhalations of LTC_4_ (1.6 mmol) or IL-25 (100 ng) or a combination of LTC_4_ and IL-25 and assessed 2 days after the last dose. The lung inflammatory infiltrate was assessed by hematoxylin and eosin staining (**A**) and eosinophils were visualized by Congo Red stain (**B**). (**C**) Gating strategy for ILC2s. (**D-F**) Number of all Thy1.2+ILC2s and subsets of KLRG1-ILC2s defined by FACS. (**G**) Total number of DCs (CD45^+^B220^-^SiglecF^-^ CD11c^+^CD11b^+^) and (**H**) percent of CD301b^+^ DCs. (**I-K**) Numbers of CD4 T cells, macrophages and CD8 T cells in the lung. Data are means ± SEM pooled from 3 independent experiments, each dot is a mouse, p values <0.05 indicated, Kruskal-Wallis ANOVA with Dunn’s correction for multiple comparisons.

**Fig. S2.**
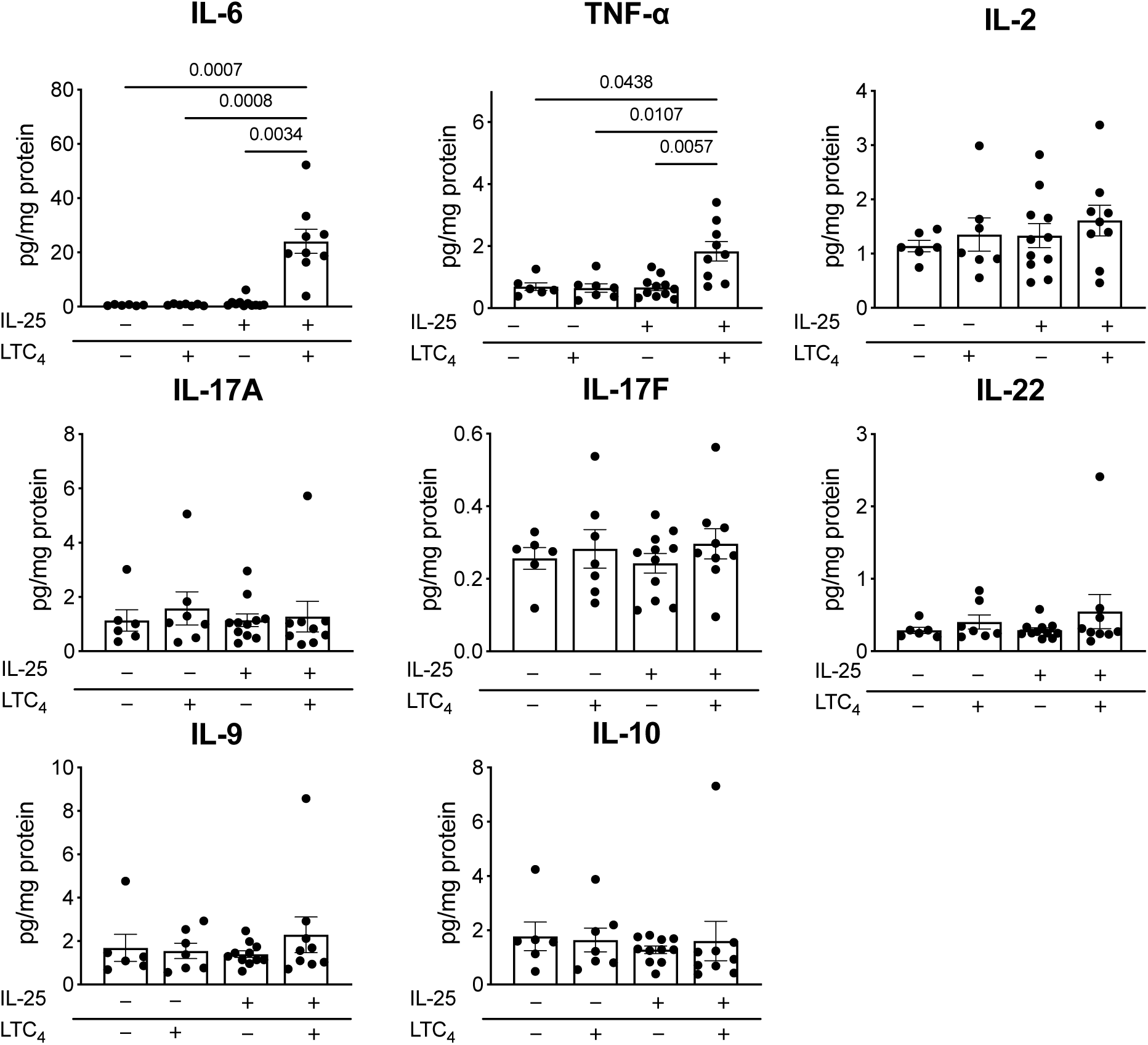
Cytokine expression profile in the lungs after intranasal challenge with LTC_4_ + IL- 25. Lung cytokine concentration was determined by LegendPlex ELISA and expressed as pg per mg of lung protein. Data are means ± SEM pooled from 3 independent experiments, each dot is a separate mouse, p values <0.05 indicated, Kruskal-Wallis ANOVA with Dunn’s correction for multiple comparisons.

**Fig. S3.**
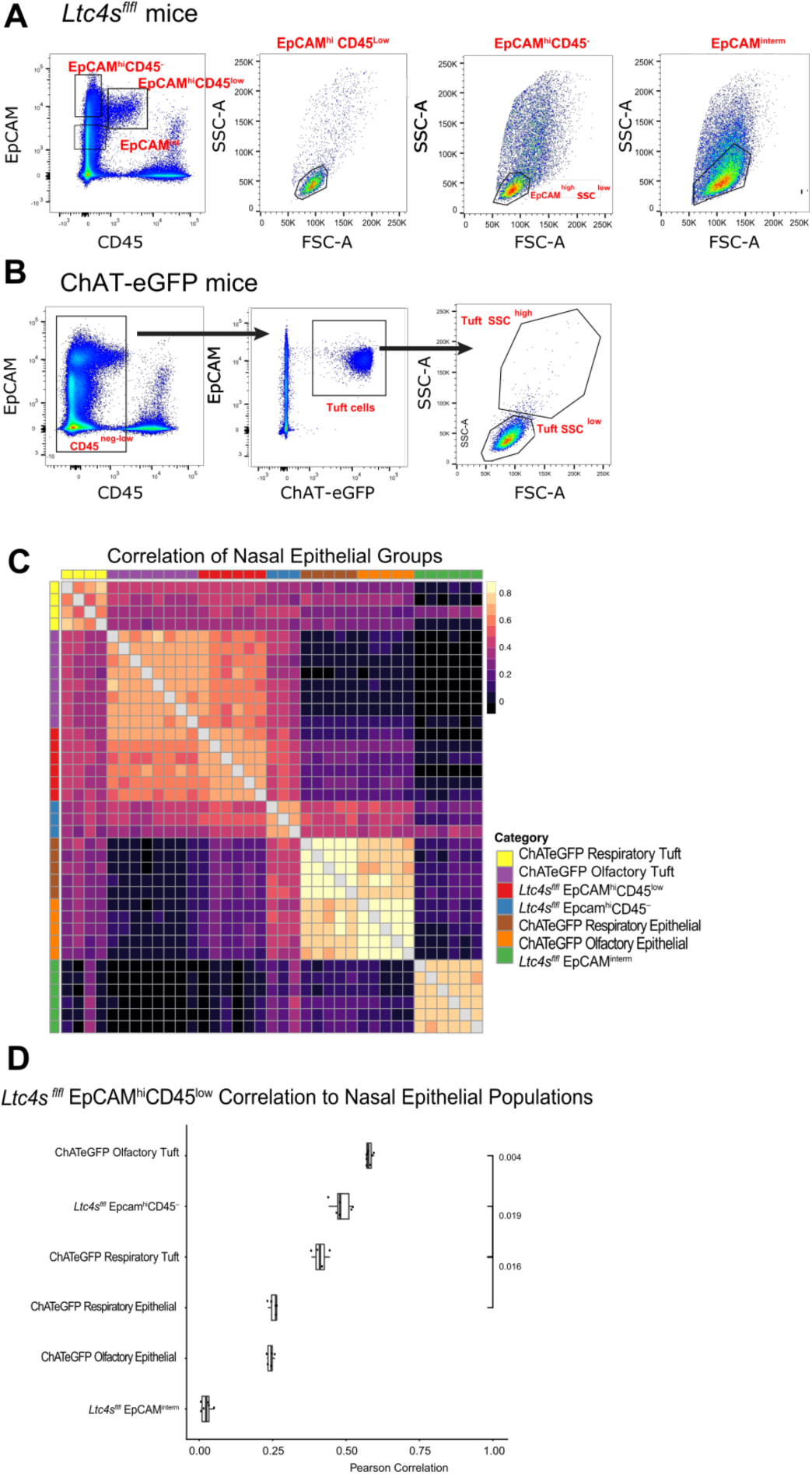
EpCAM^high^ CD45 ^low^ cells are highly enriched for tuft cells. (**A**) Gating strategy for isolation of tuft cells from *Ltc4s^fl/fl^* mice. Three populations of EpCAM^+^ cells were distinguished – EpCAM^high^-CD45^+^, EpCAM^high^-CD45^-^ and EpCAM^interm^. (**B**) Gating strategy for isolation of tuft cells from ChAT-eGFP mice. Tuft cells were defined as low or negative for CD45 and high for EpCAM. Olfactory tuft cells were defined as ChAT-eGFP^+^ cells from the olfactory mucosa with low FSC and SSC. Respiratory tuft cells were defined as tuft cells derived from the respiratory mucosa that were high for FSC and SSC. (**C**) Heatmap shows Pearson’s correlation coefficient across expression levels of all highly variable genes (Methods) between all pairs of samples from nasal epithelial groups from ChAT-eGFP and *Ltc4s^fl/fl^* mice, samples are ordered by cluster assignment (color legend). (**D**) Pearson’s correlation coefficient (x-axis) between *Ltc4s^fl/fl^* EpCAM^high^CD45^low^ tuft cell-enriched samples (points) and populations of pure tuft cells from ChAT-eGFP mice and other epithelial populations. Boxplots show the median correlation and interquartile range. P-values calculated using Mann-Whitney U-test.

**Fig. S4.**
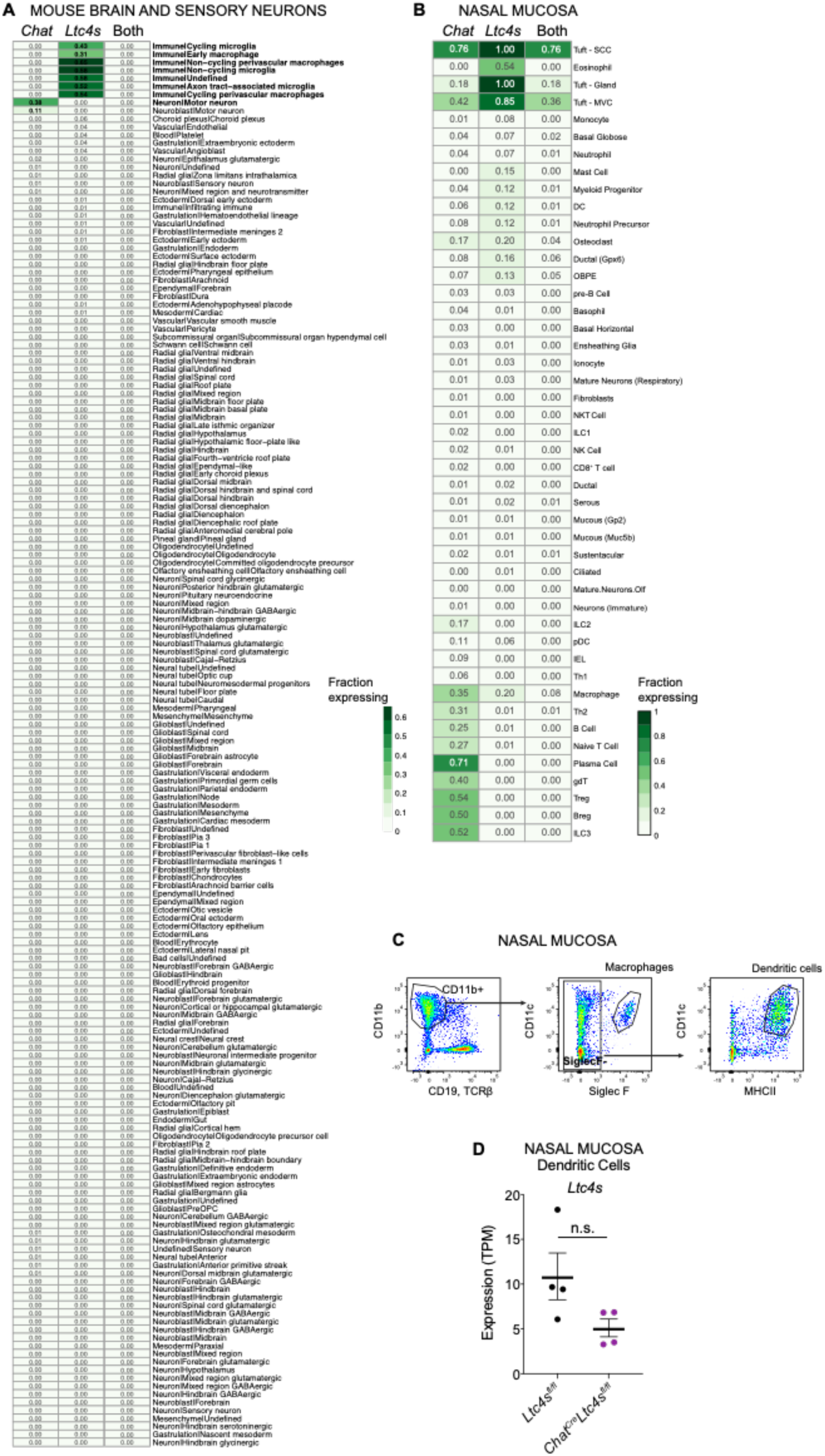
*Chat* and *Ltc4s* are specifically co-expressed in tuft cells. (**A**) Fraction of cells expressing *Chat* (first column), *Ltc4s* (second column) and both transcripts (third column) in central and peripheral neurons and nervous system immune cells from previously published data downloaded from the Mouse Brain Atlas (*36*). (**B**) Fraction of cells expressing *Chat* (first column), *Ltc4s* (second column) and both transcripts (third column). Data derived from a newly generated unpublished dataset in our lab of 50, 000 nasal epithelial and immune cells. (**C**) DCs and macrophages were defined within the CD45^+^ cell subset as CD19^−^ (B cell marker), TCRβ^−^ (T cell marker), CD11b^+^, and SiglecF^+^ for macrophages. DCs were further defined as SiglecF^−^ CD11c^+^ and MHCII^+^. (**D**) Expression level in transcripts per million (TPM) of *Ltc4s* in DCs derived from *Ltc4s^fl/fl^* and *Chat^Cre^Ltc4s^fl/fl^* mice, p adjusted value derived from DeSeq2 analysis, n.s.– not significant.

**Fig. S5.**
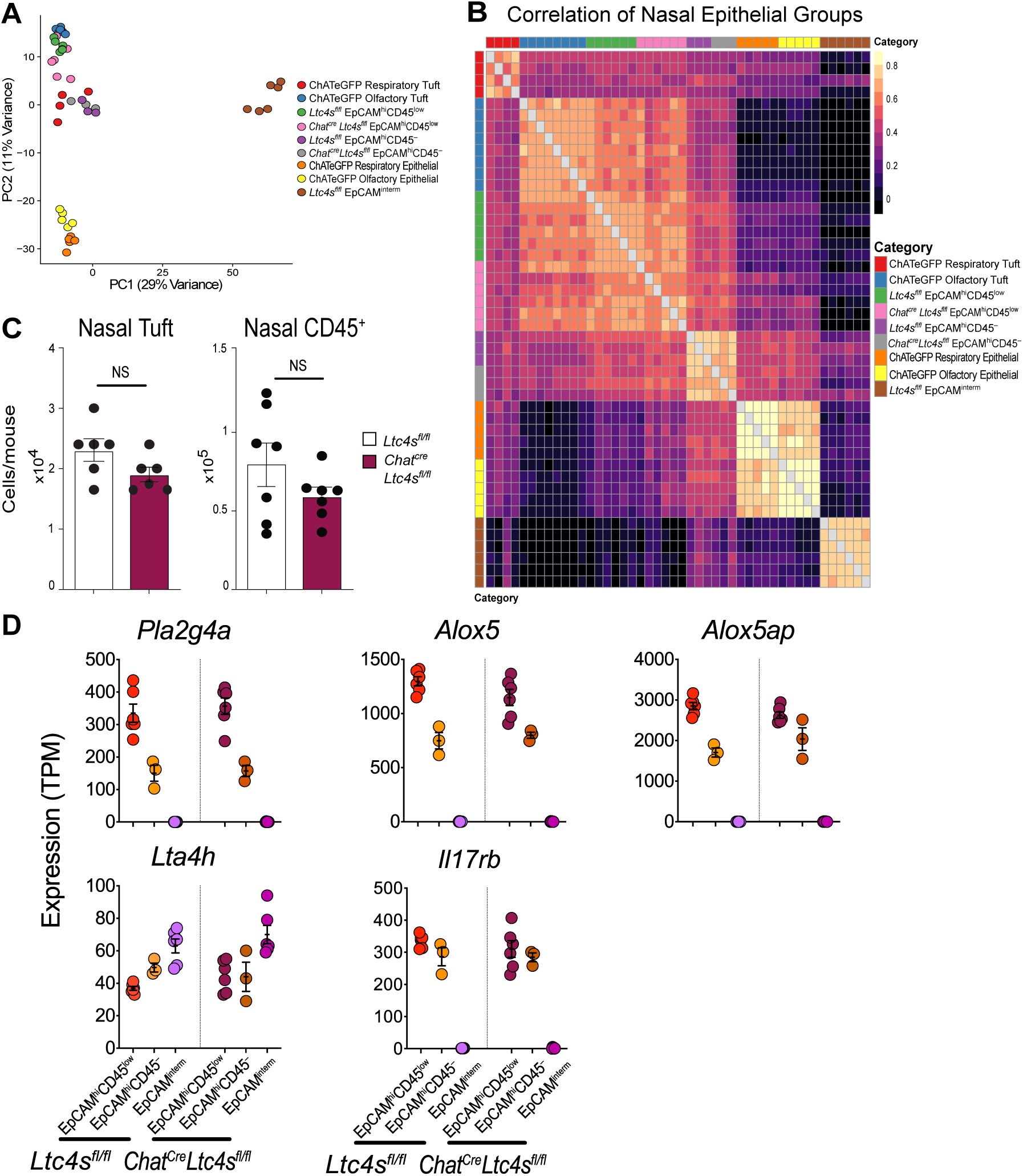
The transcriptional profile of tuft cells from *Chat^Cre^Ltc4s^fl/fl^* mice is unaltered. (**A**) Principal component analysis of tuft cells from ChAT-eGFP mice and tuft cells enriched and not enriched populations of epithelial cells derived from the nasal mucosa of ChAT-eGPF mice, *Ltc4s^fl/fl^* and *Chat^Cre^Ltc4s^fl/fl^* mice. Numbers indicate frequency of transcripts described by each principal component. (**B**) Pearson correlation coefficient (*r*) between the nasal epithelial groups from ChAT-eGFP, *Ltc4s^fl/fl^* and *Chat^Cre^Ltc4s^fl/fl^* mice ordered by cluster assignment. (**C**) Numbers of nasal tuft cells (EpCAM^high^CD45^low^SSC^low^) and nasal CD45^+^ cells in *Ltc4s^fl/fl^* and *Chat^Cre^Ltc4s^fl/fl^* mice derived from cell sorting. NS = not significant.(**D**) Expression level in transcripts per million (TPM) of the indicated genes in epithelial cell subsets from *Ltc4s^fl/fl^* and *Chat^Cre^Ltc4s^fl/fl^* mice.

**Fig. S6.**
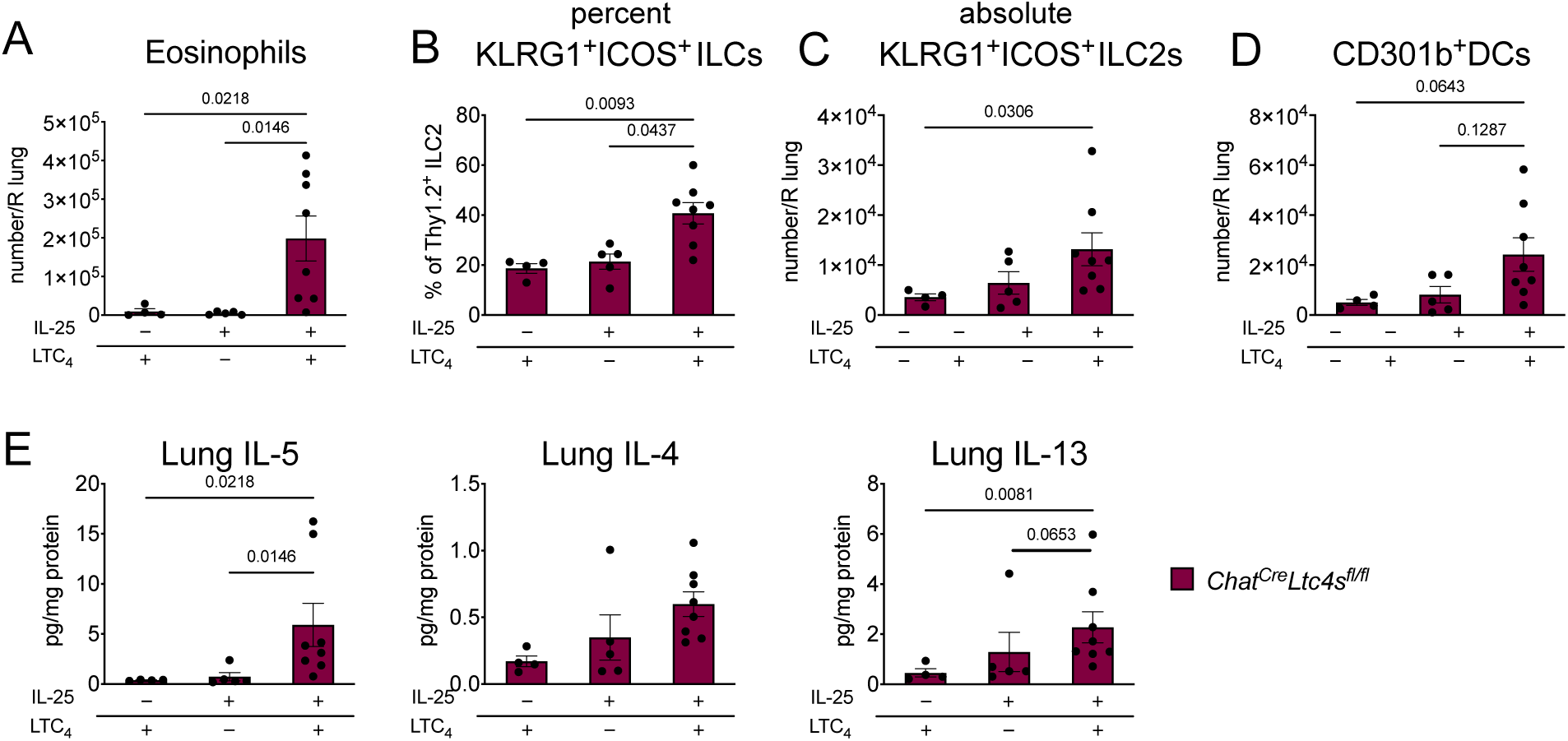
The synergy of IL-25 and LTC4 is preserved in *Chat^Cre^Ltc4s^fl/fl^* mice. *Chat^Cre^Ltc4s^fl/fl^* mice were given LTC_4_ (1.6 mmol), or IL-25 (100 ng) or a combination of LTC_4_ and IL-25 and assessed 2 days after the last dose. The frequency of eosinophils (**a**), frequency and number of KLRG1^+^ICOS^+^Thy1.2^+^ ILC2s (**B, C**), and number of CD301b^+^MHCII^+^CD11b^+^ DCs (**D**) were assessed by FACS. (**E**). Lung cytokine protein concentration was determined by LegendPlex. Data are means ± SEM pooled from 3 independent experiments, each dot is a mouse, p values <0.05 indicated, Kruskal-Wallis ANOVA with Dunn’s correction for multiple comparisons.

**Fig. S7.**
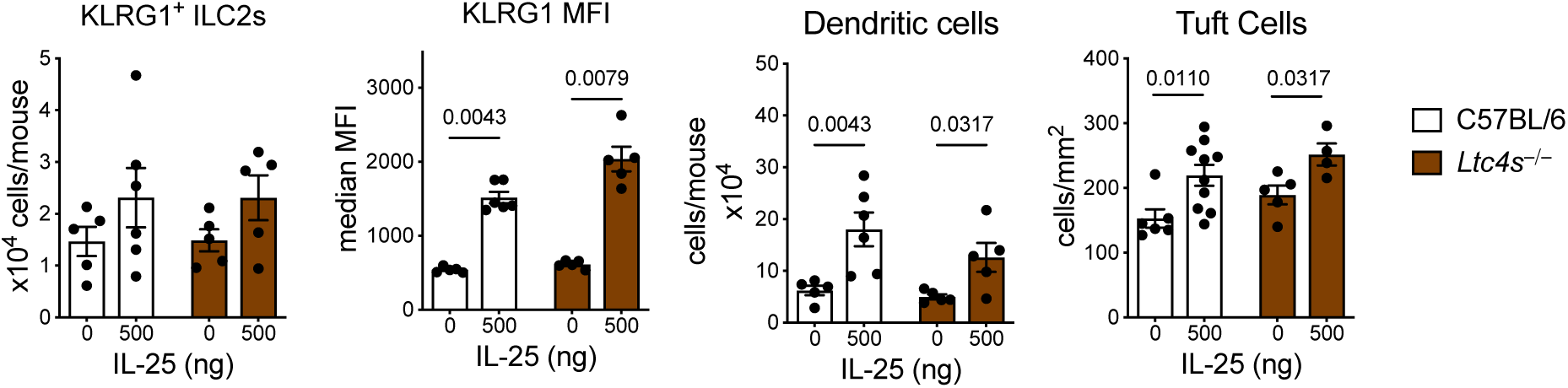
High dose IL-25 induced inflammation is preserved in *Ltc4s^-/-^* mice. WT and *Ltc4s^−/−^* mice were given 3 daily intranasal doses of 500 ng of IL-25 and assessed 2 days after the last dose. Lung KLRG1^+^ ILC2 numbers and KLRG1 MFI and DCs were assessed in the lung and tuft cells in the trachea 48 hours after the last IL-25 inhalation. Data are means + SEM from ≥ 2 experiments; each circle represents a separate mouse, Mann Whitney U test, p values < 0.05 are indicated.

## FUNDING

This work was supported by:

National Institutes of Health grant K08 AI132723 (LGB)

National Institutes of Health grant 1R21AI154345 (LGB and ALH)

National Institutes of Health grant R01AI078908

National Institutes of Health grant R37AI052353 (JAB)

R01AI136041 (JAB)

R01HL136209 (JAB)

National Institutes of Health grant R01 AI134989 (NAB) U19 AI095219 (NAB, JAB)

AAAAI Foundation Faculty Development Award (LGB)

Joycelyn C. Austen Fund for Career Development of Women Physician Scientists (LGB) generous donation by the Vinik family (LGB)

## Author contributions

Conceptualization: LGB, ALH and JAB

Methodology: LGB, ALH, EL, SU

Investigation: LGB, SU, AAB, CW, EL, JL

Writing: LGB, AHL, JAB, NAB, IM

Funding Acquisition: LGB, JAB, NAB, AHL

Resources: TL, IM and JAB

## Acknowledgements

We thank Adam Chicoine from Brigham and Women’s Hospital Human Immunology Center Flow Core for his help with FACS sorting. We are grateful to Charles Vidoudez and Sunia Trauger from the Harvard Center for Mass Spectrometry and Oswald Quehenberger from the University of California San Diego Lipidomics Core for their expertise and help setting up the mass spectrometry assays for CysLTs and lipidomics assays. We are also thankful to Lisa Goodrich at the Department of Neurobiology at Harvard Medical School for providing reagents and infrastructure for in situ hybridization. Schematic diagrams were created using BioRender.

