## SupplementalFile2 for "Tuft cell-produced cysteinyl leukotrienes and IL-25 synergistically initiate lung type 2 inflammation"

### SUPPLEMENTARY MATERIALS:

**Fig. S1. $LTC_4$ and IL-25 synergize for airway type 2 lung inflammation.**
**Fig. S2. Cytokine expression profile in the lungs after intranasal challenge with $LTC_4$ + IL-25.**
**Fig. S3. EpCAM$^{high}$ CD45 $^{low}$ cells are highly enriched for tuft cells.**
**Fig. S4. *Chat* and *Ltc4s* are specifically co-expressed in tuft cells.**
**Fig. S5. The transcriptional profile of tuft cells from *Chat$^{Cre}$Ltc4s$^{fl/fl}$* mice is unaltered.**
**Fig. S6. The synergy of IL-25 and LTC4 is preserved in *Chat$^{Cre}$Ltc4s$^{fl/fl}$* mice.**
**Fig. S7. High dose IL-25 induced inflammation is preserved in *Ltc4s$^{-/-}$* mice.**
**Table 1. Materials**

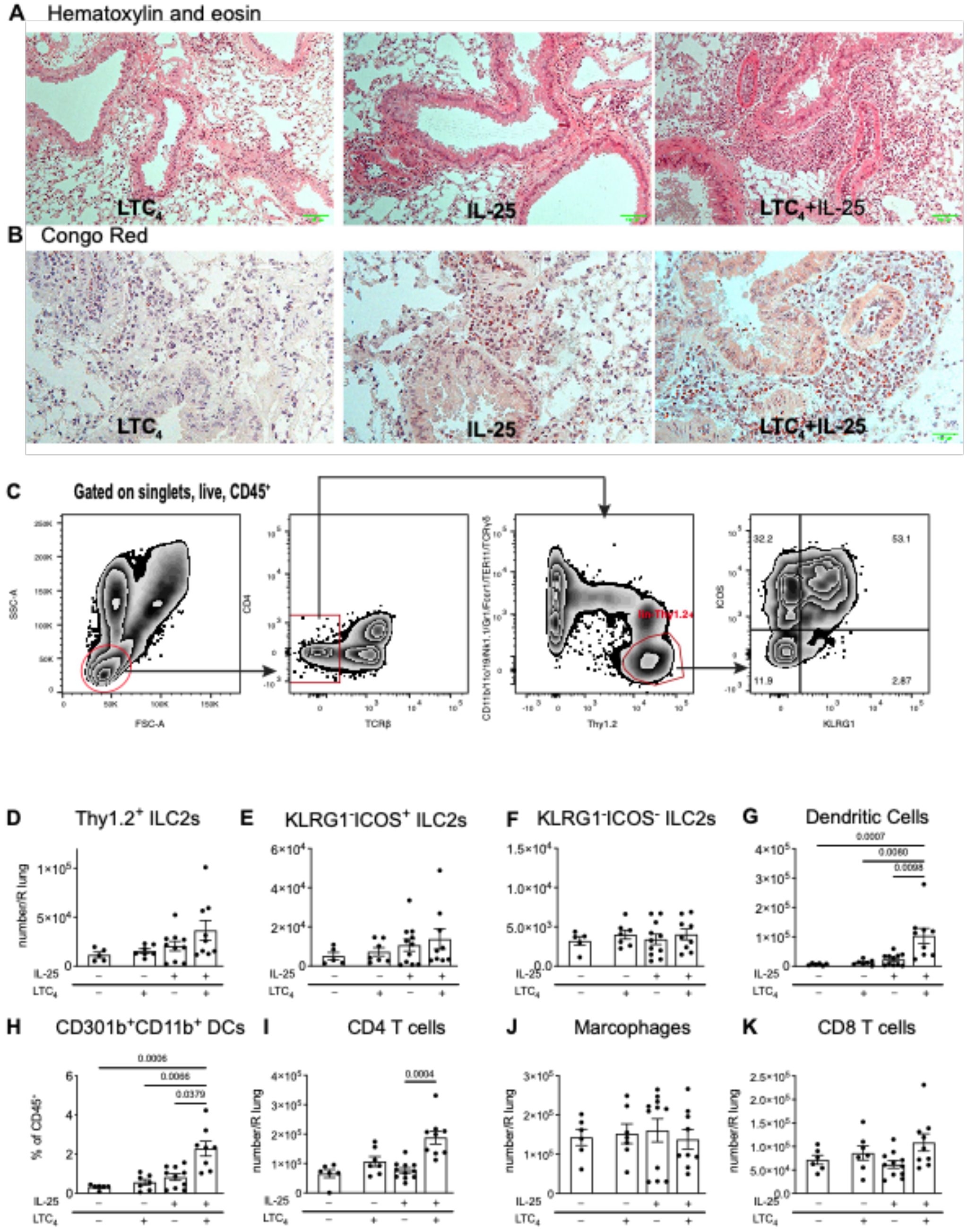

**Fig. S1. $LTC_4$ and IL-25 synergize for airway type 2 lung inflammation.**
(**A**) WT (C57BL/6 and *Ltc4s*$^{fl/fl}$) or *Chat*$^{Cre}$*Ltc4s*$^{fl/fl}$ mice were given three daily inhalations of $LTC_4$ (1.6 mmol) or IL-25 (100 ng) or a combination of $LTC_4$ and IL-25 and assessed 2 days after the last dose. The lung inflammatory infiltrate was assessed by hematoxylin and eosin staining (**A**) and eosinophils were visualized by Congo Red stain (**B**). (**C**) Gating strategy for ILC2s. (**D-F**) Number of all Thy1.2+ILC2s and subsets of KLRG1- ILC2s defined by FACS. (**G**) Total number of DCs ($CD45^+B220^-SiglecF^-CD11c^+CD11b^+$) and (**H**) percent of $CD301b^+$ DCs. (**I-K**) Numbers of CD4 T cells, macrophages and CD8 T cells in the lung. Data are means ± SEM pooled from 3 independent experiments, each dot is a mouse, p values <0.05 indicated, Kruskal-Wallis ANOVA with Dunn's correction for multiple comparisons.

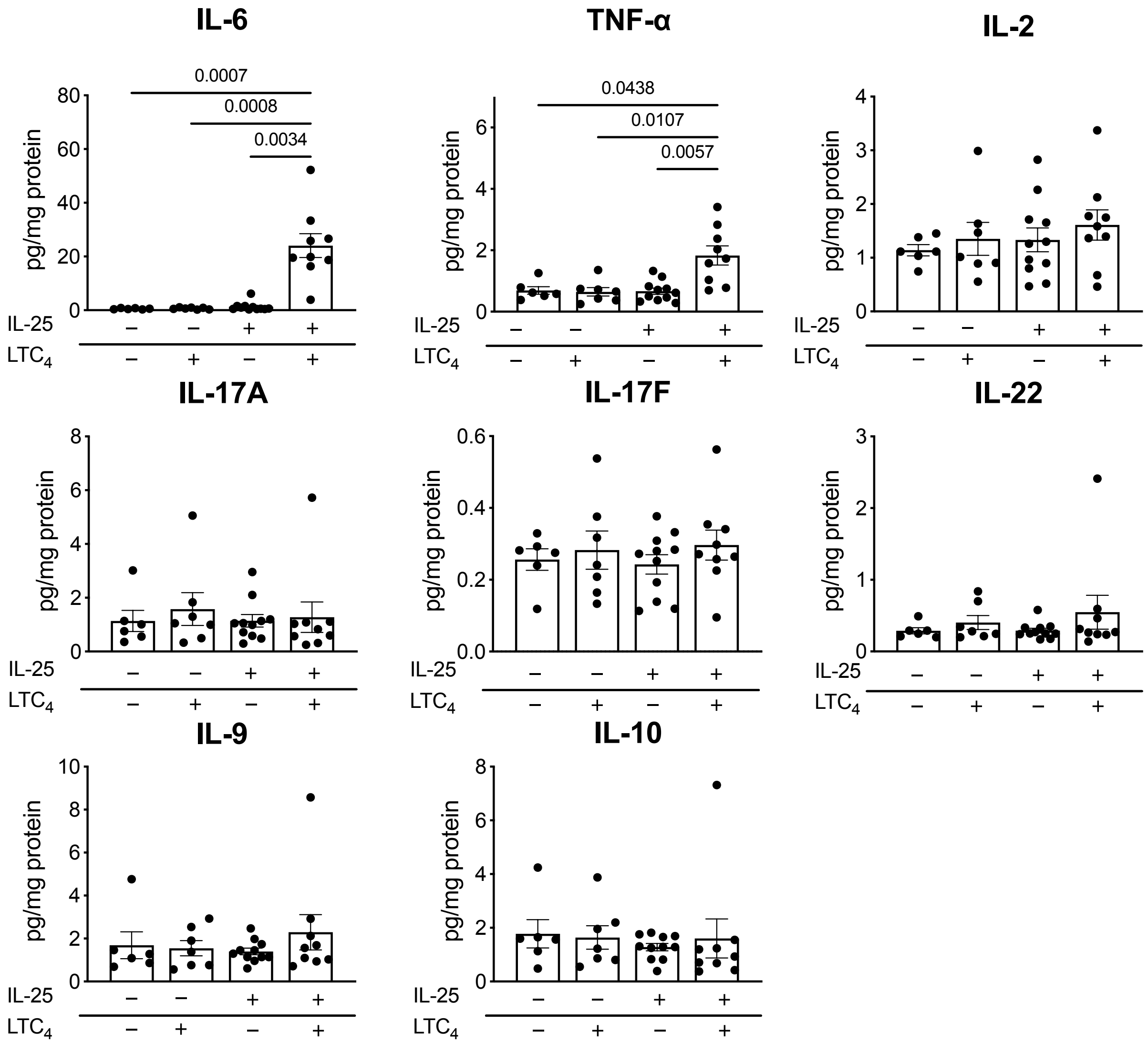

**Fig. S2. Cytokine expression profile in the lungs after intranasal challenge with $LTC_4$ + IL-25.**

Lung cytokine concentration was determined by LegendPlex ELISA and expressed as pg per mg of lung protein. Data are means ± SEM pooled from 3 independent experiments, each dot is a separate mouse, p values <0.05 indicated, Kruskal-Wallis ANOVA with Dunn’s correction for multiple comparisons.

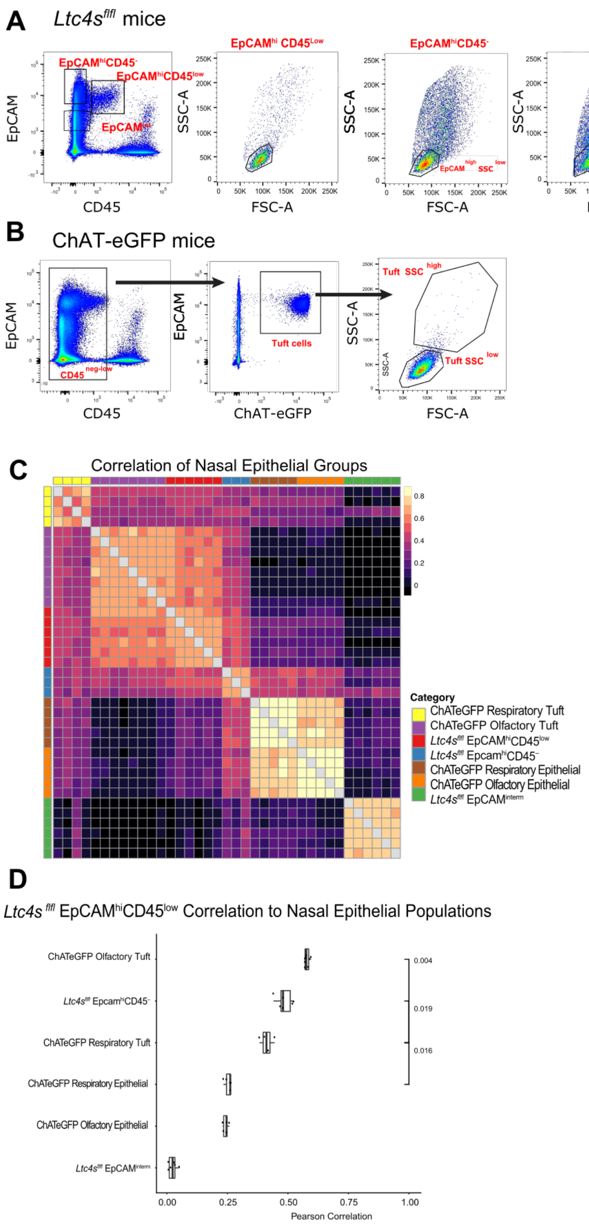

**Fig. S3. EpCAM$^{high}$ CD45 $^{low}$ cells are highly enriched for tuft cells.** (**A**) Gating strategy for isolation of tuft cells from *Ltc4s$^{fl/fl}$* mice. Three populations of EpCAM$^{+}$ cells were distinguished – EpCAM$^{high}$-CD45$^{+}$, EpCAM$^{high}$-CD45$^{-}$ and EpCAM$^{interm}$. (**B**) Gating strategy for isolation of tuft cells from ChAT-eGFP mice. Tuft cells were defined as low or negative for CD45 and high for EpCAM. Olfactory tuft cells were defined as ChAT-eGFP$^{+}$ cells from the olfactory mucosa with low FSC and SSC. Respiratory tuft cells were defined as tuft cells derived from the respiratory mucosa that were high for FSC and SSC. (**C**) Heatmap shows Pearson's correlation coefficient across expression levels of all highly variable genes (Methods) between all pairs of samples from nasal epithelial groups from ChAT-eGFP and *Ltc4s$^{fl/fl}$* mice, samples are ordered by cluster assignment (color legend). (**D**) Pearson's correlation coefficient (x-axis) between *Ltc4s$^{fl/fl}$* EpCAM$^{high}$CD45$^{low}$ tuft cell-enriched samples (points) and populations of pure tuft cells from ChAT-eGFP mice and other epithelial populations. Boxplots show the median correlation and interquartile range. P-values calculated using Mann-Whitney U-test.

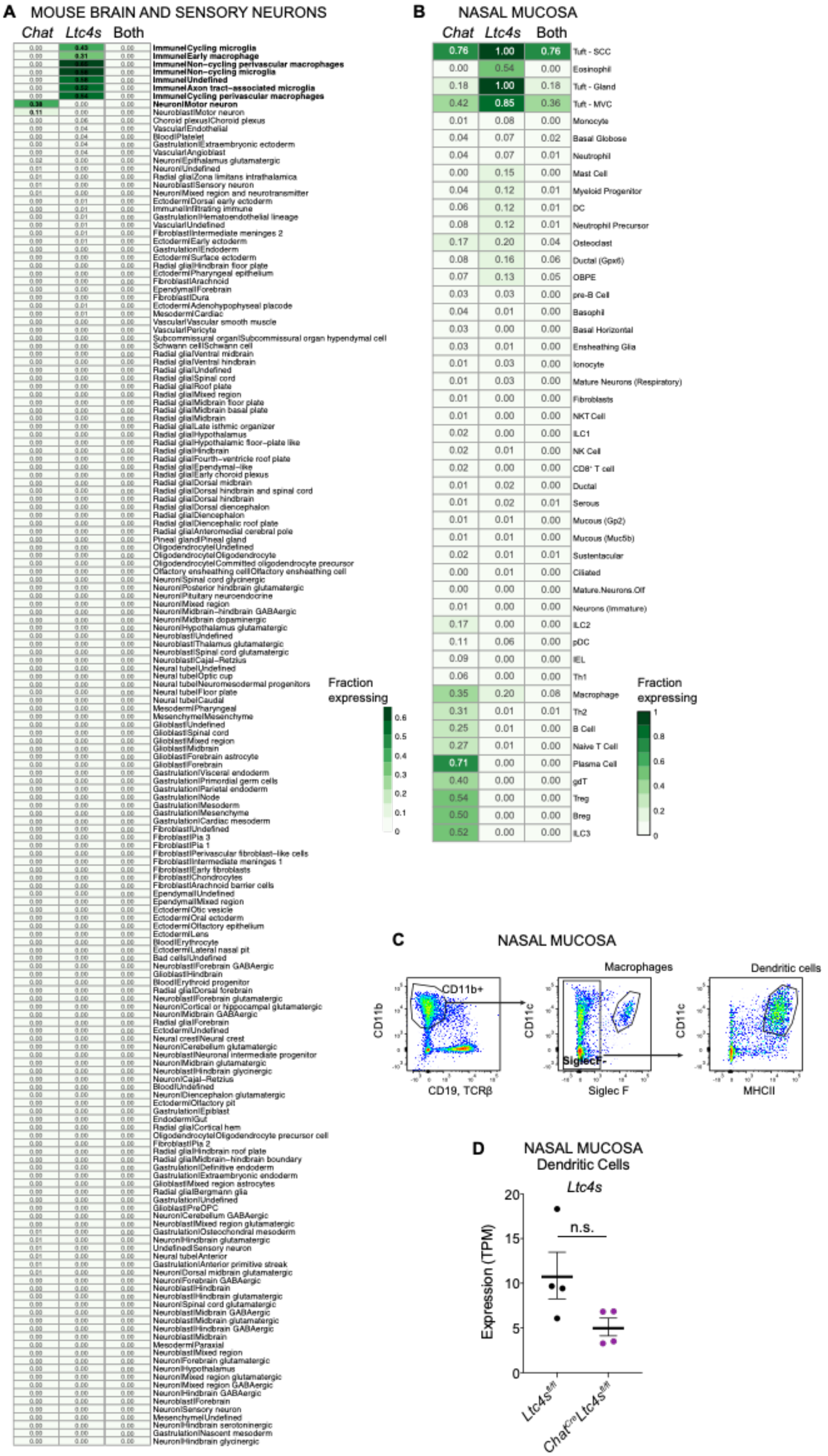
A
MOUSE BRAIN AND SENSORY NEURONS
Chat Ltc4s Both
Fraction expressing
B
NASAL MUCOSA
Chat Ltc4s Both
0.76 1.00 0.76 Tuft - SCC
0.00 0.54 0.00 Eosinophil
0.18 1.00 0.18 Tuft - Gland
0.42 0.85 0.36 Tuft - MVC
0.01 0.08 0.00 Monocyte
0.04 0.07 0.02 Basal Globose
0.04 0.07 0.01 Neutrophil
0.00 0.15 0.00 Mast Cell
0.04 0.12 0.01 Myeloid Progenitor
0.06 0.12 0.01 DC
0.08 0.12 0.01 Neutrophil Precursor
0.17 0.20 0.04 Osteoclast
0.08 0.16 0.06 Ductal (Gpx6)
0.07 0.13 0.05 OBPE
0.03 0.03 0.00 pre-B Cell
0.04 0.01 0.00 Basophil
0.03 0.00 0.00 Basal Horizontal
0.03 0.01 0.00 Ensheathing Glia
0.01 0.03 0.00 Ionocyte
0.01 0.03 0.00 Mature Neurons (Respiratory)
0.01 0.00 0.00 Fibroblasts
0.01 0.01 0.00 NKT Cell
0.02 0.00 0.00 ILC1
0.02 0.01 0.00 NK Cell
0.02 0.00 0.00 CD8+ T cell
0.01 0.02 0.00 Ductal
0.01 0.02 0.01 Serous
0.01 0.01 0.00 Mucous (Gp2)
0.01 0.01 0.00 Mucous (Muc5b)
0.02 0.01 0.01 Sustentacular
0.00 0.01 0.00 Ciliated
0.00 0.00 0.00 Mature.Neurons.Olf
0.01 0.00 0.00 Neurons (Immature)
0.17 0.00 0.00 ILC2
0.11 0.06 0.00 pDC
0.09 0.00 0.00 IEL
0.06 0.00 0.00 Th1
0.35 0.20 0.08 Macrophage
0.31 0.01 0.01 Th2
0.25 0.01 0.00 B Cell
0.27 0.01 0.00 Naive T Cell
0.71 0.00 0.00 Plasma Cell
0.40 0.00 0.00 gdT
0.54 0.00 0.00 Treg
0.50 0.00 0.00 Breg
0.52 0.00 0.00 ILC3
Fraction expressing
C
NASAL MUCOSA
CD11b+
CD11b
CD19, TCRβ
Macrophages
CD11c
Siglec F
SiglecF-
Dendritic cells
CD11c
MHCII
D
NASAL MUCOSA
Dendritic Cells
Ltc4s
n.s.
Expression (TPM)
Ltc4s fl/fl
ChatCre Ltc4s fl/fl

**Fig. S4. *Chat* and *Ltc4s* are specifically co-expressed in tuft cells.** (**A**) Fraction of cells expressing *Chat* (first column), *Ltc4s* (second column) and both transcripts (third column) in central and peripheral neurons and nervous system immune cells from previously published data downloaded from the Mouse Brain Atlas *(36)*. (**B**) Fraction of cells expressing *Chat* (first column), *Ltc4s* (second column) and both transcripts (third column). Data derived from a newly generated unpublished dataset in our lab of 50, 000 nasal epithelial and immune cells. (**C**) DCs and macrophages were defined within the $CD45^{+}$ cell subset as $CD19^{-}$ (B cell marker), $TCR\beta^{-}$ (T cell marker), $CD11b^{+}$, and $SiglecF^{+}$ for macrophages. DCs were further defined as $SiglecF^{-}$ $CD11c^{+}$ and $MHCII^{+}$. (**D**) Expression level in transcripts per million (TPM) of *Ltc4s* in DCs derived from $Ltc4s^{fl/fl}$ and $Chat^{Cre}Ltc4s^{fl/fl}$ mice, p adjusted value derived from DeSeq2 analysis, n.s.– not significant.

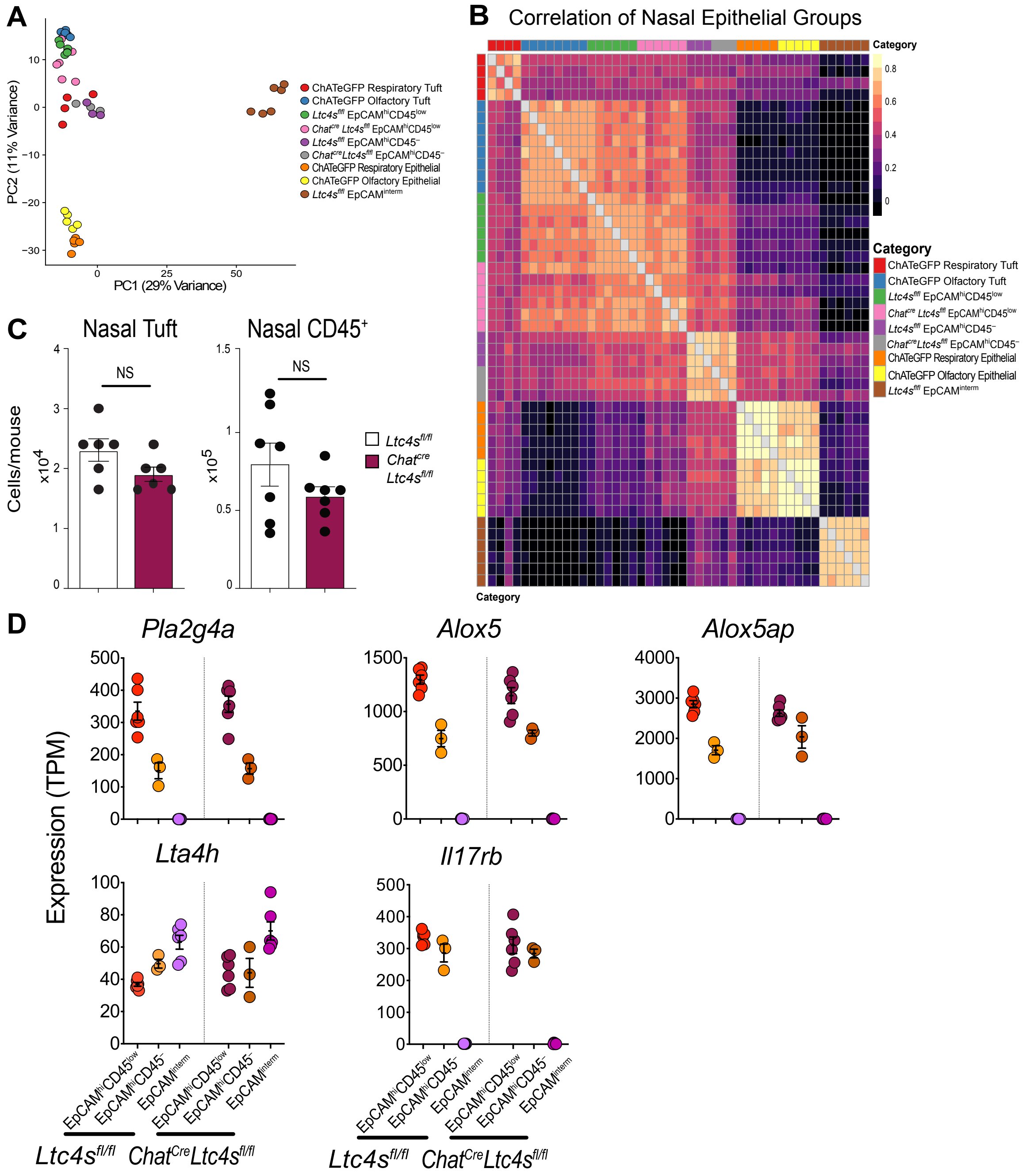

**Fig. S5. The transcriptional profile of tuft cells from *Chat*$^{Cre}$*Ltc4s*$^{fl/fl}$ mice is unaltered.** (**A**) Principal component analysis of tuft cells from ChAT-eGFP mice and tuft cells enriched and not enriched populations of epithelial cells derived from the nasal mucosa of ChAT-eGPF mice, *Ltc4s*$^{fl/fl}$ and *Chat*$^{Cre}$*Ltc4s*$^{fl/fl}$ mice. Numbers indicate frequency of transcripts described by each principal component. (**B**) Pearson correlation coefficient (*r*) between the nasal epithelial groups from ChAT-eGFP, *Ltc4s*$^{fl/fl}$ and *Chat*$^{Cre}$*Ltc4s*$^{fl/fl}$ mice ordered by cluster assignment. (**C**) Numbers of nasal tuft cells (EpCAM$^{high}$CD45$^{low}$SSC$^{low}$) and nasal CD45$^{+}$ cells in *Ltc4s*$^{fl/fl}$ and *Chat*$^{Cre}$*Ltc4s*$^{fl/fl}$ mice derived from cell sorting. NS = not significant.(**D**) Expression level in transcripts per million (TPM) of the indicated genes in epithelial cell subsets from *Ltc4s*$^{fl/fl}$ and *Chat*$^{Cre}$*Ltc4s*$^{fl/fl}$ mice.

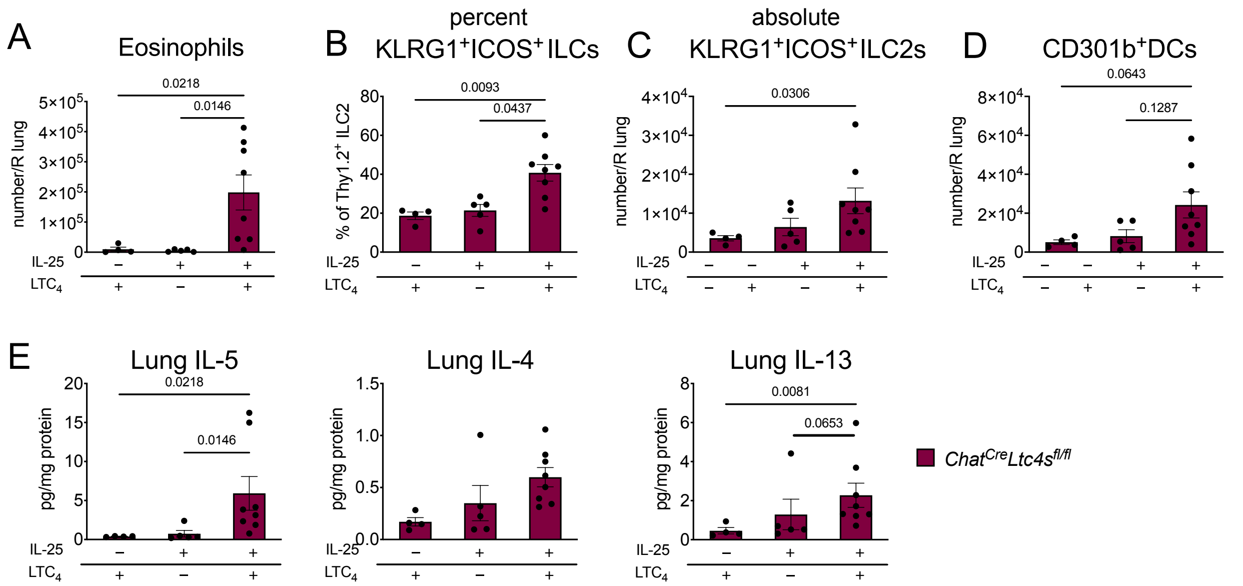

**Fig. S6. The synergy of IL-25 and LTC4 is preserved in *Chat*$^{Cre}$*Ltc4s*$^{fl/fl}$ mice.** *Chat*$^{Cre}$*Ltc4s*$^{fl/fl}$ mice were given $LTC_4$ (1.6 mmol)’ or IL-25 (100 ng) or a combination of $LTC_4$ and IL-25 and assessed 2 days after the last dose. The frequency of eosinophils (**a**), frequency and number of KLRG1$^+$ICOS$^+$Thy1.2$^+$ ILC2s (**B, C**), and number of CD301b$^+$MHCII$^+$CD11b$^+$ DCs (**D**) were assessed by FACS. (**E**). Lung cytokine protein concentration was determined by LegendPlex. Data are means ± SEM pooled from 3 independent experiments, each dot is a mouse, p values <0.05 indicated, Kruskal-Wallis ANOVA with Dunn’s correction for multiple comparisons.

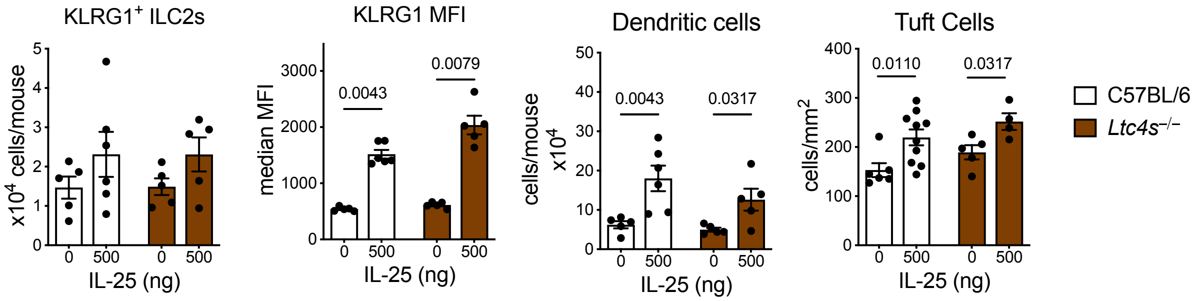

**Fig. S7. High dose IL-25 induced inflammation is preserved in *Ltc4s*$^{-/-}$ mice.** WT and *Ltc4s*$^{-/-}$ mice were given 3 daily intranasal doses of 500 ng of IL-25 and assessed 2 days after the last dose. Lung KLRG1$^+$ ILC2 numbers and KLRG1 MFI and DCs were assessed in the lung and tuft cells in the trachea 48 hours after the last IL-25 inhalation. Data are means + SEM from ≥ 2 experiments; each circle represents a separate mouse, Mann Whitney U test, p values < 0.05 are indicated.

**TABLE 1**

**Name of Material/ Equipment** **Company** **Catalog Number**

Reagents

| REAGENT or RESOURCE | SOURCE | IDENTIFIER |
|---|---|---|
| Antibodies | | |
| Anti mouse/human DCLK1 (Polyclonal Rabbit Ig) | Abcam | cat# ab31704 |
| Normal Donkey Serum | Jackson ImmunoResearch | cat# 017-000-121 |
| donkey anti-rabbit IgG (H+L) Highly Cross-Adsorbed Secondary Antibody, Alexa Fluor 594 | Life Technologies/Molecular Probes | cat#A-21207 |
| Trustain FC Block | Biolegend | cat# 101320 |
| APC anti-mouse CD326 (EpCAM) | Biolegend | cat#118214 |
| Brilliant Violet 421 anti-mouse CD45 | Biolegend | cat#103134 |
| FITC anti-mouse I-A/I-E | Biolegend | cat#107606 |
| Brilliant Violet 421 anti-mouse CD8a | Biolegend | cat#100748 |
| APC/Cy7 anti-mouse/human CD11b | Biolegend | cat#101226 |
| FITC anti-mouse FcεRIα | Biolegend | cat#134306 |
| FITC anti-mouse CD11c | Biolegend | cat#117306 |
| PerCP/Cy5.5 anti-mouse Siglec F | BD Biosciences | cat# 565526 |
| FITC Rat anti-mouse IgE | BD Biosciences | cat#553415 |

| Chemicals, Peptides, and Recombinant Proteins | | |
|---|---|---|
| Alternaria alternata culture filtrate (lot# 151774) | Greer Laboratories | cat#XPM1C3A25 |
| *Dermatophagoides pteronyssinus* extract (lot# 45415) | Greer Laboratories | cat#B82 |
| Hoechst 33342 | Sigma | cat#14533 |
| DNAse I | Sigma | cat# 10104159001 |
| Dispase | Gibco | cat# 17105041 |

| | | |
|---|---|---|
| HEPES-Tyrode's Buffer Without Calcium | Boston BioProducts | cat# PY-912 |
| Tyrode's Solution (HEPES-Buffered) | Boston BioProducts | cat# BSS-355 |
| L-Cysteine | Sigma | cat# C7352 |
| Papain from papaya latex | Sigma | cat# P3125 |
| Leupeptin trifluoroacetate salt | Sigma | cat# L2023 |
| Propidium iodide | Sigma | cat# P4170 |
| A23187 | Cayman | cat# 11016 |
| ATP-γ-S | Abcam | cat# ab138911 |
| 2-Mercaptoethanol | Gibco | cat#21985023 |
| Cysteinyl Leukotriene ELISA Kit | Cayman | cat# 500390 |
| Prostaglandin D2 ELISA Kit | Cayman | cat# 512031 |
| Target Retrieval Solution 10x Concentrate | Dako | cat# S1699 |
| Leukotriene C4 | Cayman Chemical | cat#20210 |
| N-methyl Leukotriene C4 | Cayman Chemical | cat#13390 |
| Recombinant mouse IL17PE/IL-25 | R&D systems | cat#7909-IL/CF |
| LEGENDplex™ MU Th Cytokine Panel (12-plex) w/ VbP V03 | Biolegend | cat#741044 |
| Pierce T-PER™, Tissue Protein Extraction Reagent | ThermoFisher | cat#78510 |
| Roche cOmplete™ Protease Inhibitor Cocktail | Sigma Aldrich | cat#4693116001 |
| BCA assay | ThermoFisher | cat# 23225 |
| IL-33 ELISA | Invitrogen | cat# 88733322 |
| TCL buffer | Qiagen | cat# 1031576 |
| β-mercaptoethanol | Sigma | cat #M6250 |

| Media | | |
|---|---|---|
| DMEM/F-12 | ThermoFischer | cat#11320033 |
| HEPES | ThermoFischer | cat#AAJ1692622 |
| Penicillin-Streptomycin | ThermoFischer | cat#15140148 |
| Fetal Bovine Serum | ThermoFischer | cat#10082139 |
| Nu-serum | Corning | cat#355100 |

| | | |
|---|---|---|
| L-glutamin | | |
| BSA | ThermoFischer | cat#BP9706100 |
| Amphotericin B | Gibco | cat#1529001 |
| RPMI | Corning | cat#10040CV |

| Experimental Models: Organisms/Strains | | |
|---|---|---|
| C57BL/6 Mouse | Charles River | 27 |
| *Ltc4s*$^{-/-}$ | Dr. Yoshihide Kanaoka | |
| *ChAT*$^{BAC}$*-eGFP (B6.Cg-Tg(RP23-268L19-EGFP)2Mik/J)* | The Jackson Laboratory | 7902 |
| *ChAT-IRES-Cre::frt-neo-frt (Chat*$^{tm2(cre)Lowl}$*)/J* | The Jackson Laboratory | 006410 |
| *Ltc4S*$^{fl/fl}$ | Dr. Joshua Boyce | |
| Mcpt5/DTA | Axel Roers | |
| Mcpt5-Cre | Axel Roers | |
| *Cysltr2*$^{-/-}$ | Dr. Yoshihide Kanaoka | |
| *Oxgr1-/-* | Dr. Yoshihide Kanaoka | |
| C57BL/6N-Cysltr1tm1$^{Ykn}$/J | The Jackson Laboratory | 30814 |
| *Pou2f3-/-* | Ichiro Matsumoto | |

| Software and Algorithms | | |
|---|---|---|
| FlowJo v.8 | FlowJo, LLC | https://www.www.flowjo.com |
| Prism 8 | GraphPad Software | https://www.graphpad.com/scientific-software/prism/ |

| | | |
|---|---|---|
| R and R studio | | http://www.r-project.com/ |
| ImageJ | | National Institutes of Health, Bethesda, MD |
